# Chromatin-based techniques map DNA interaction landscapes in psoriasis susceptibility loci and highlight *KLF4* as a target gene in 9q31

**DOI:** 10.1101/822304

**Authors:** Helen Ray-Jones, Kate Duffus, Amanda McGovern, Paul Martin, Chenfu Shi, Jenny Hankinson, Oliver Gough, Annie Yarwood, Andrew P Morris, Antony Adamson, Christopher Taylor, James Ding, Vasanthi Priyadarshini Gaddi, Yao Fu, Patrick Gaffney, Gisela Orozco, Richard B Warren, Steve Eyre

**Affiliations:** Centre for Genetics and Genomics Versus Arthritis. Division of Musculoskeletal and Dermatological Sciences, School of Biological Sciences, Faculty of Biology, Medicine and Health, The University of Manchester, United Kingdom; Dermatology Centre, Salford Royal NHS Foundation Trust, Manchester NIHR Biomedical Research Centre, Manchester Academic Health Science Centre, Manchester, United Kingdom; Genes and Human Disease Research Program, Oklahoma Medical Research Foundation, Oklahoma City, 73104, OK, USA; The Lydia Becker Institute of Immunology and Inflammation, University of Manchester, Manchester, UK; Genome Editing Unit, Faculty of Biology, Medicine and Health, The University of Manchester, Manchester, United Kingdom

## Abstract

Genome-wide association studies (GWAS) have uncovered many genetic risk loci for psoriasis, yet many remain uncharacterised in terms of the causal gene and their biological mechanism in disease. Here, we use a disease-focused Capture Hi-C experiment to link psoriasis-associated variants with their target genes in psoriasis-relevant cell lines (HaCaT keratinocytes and My-La CD8+ T cells). We confirm previously assigned genes, suggest novel candidates and provide evidence for complexity at psoriasis GWAS loci. In the 9q31 risk locus we combine further epigenomic evidence to demonstrate how the psoriasis association forms a functional interaction with the distant (>500 kb) *KLF4* gene. We use CRISPR activation coupled with RNA-seq to demonstrate how activation of psoriasis-associated enhancers upregulates *KLF4* in HaCaT cells. Our study design provides a robust pipeline for following up on GWAS disease-associated variants, paving the way for functional translation of genetic findings into clinical benefit.

## Introduction

Psoriasis is an immune-mediated condition affecting around 2% of the worldwide population (Griffiths and Barker, 2007). Psoriasis usually manifests as red, scaly plaques on the skin and is thought to be driven by a number of immune pathways; predominantly T helper cell Th1 and Th17 signalling. Although lifestyle factors play a role in disease susceptibility, genetic predisposition is the largest risk factor for psoriasis. Genome-wide association studies (GWAS) have so far identified 63 loci associated with susceptibility in individuals of European ancestry (Tsoi et al., 2017) and more than 20 further unique signals in the Han Chinese population (Yin et al., 2015, Zuo et al., 2015). The majority of single nucleotide polymorphisms (SNPs) associated with psoriasis and other immune-mediated conditions do not map within gene coding regions; rather they are enriched in non-coding enhancer elements (Kundaje et al., 2015), with approximately 60% of predicted causal SNPs lying within cell type specific immune enhancers relevant to the disease of interest, and approximately 8% in promoters (Ernst et al., 2011, Farh et al., 2015). Historically, gene candidates were assigned to GWAS loci based on proximity or biological function; however, this can lead to incorrect interpretation of results since it is now well established that enhancers can regulate genes over very large genomic distances through chromatin looping (Javierre et al., 2016, Rao et al., 2014).

The challenge now is to link disease-associated enhancers with the true genes that they regulate in order to determine the relevant cell types and the mechanism of regulation. Advances in sequencing, molecular biology and genome editing are now enabling us to answer these pivotal, ‘post-GWAS’ questions (Ray-Jones et al., 2016). Hi-C is a technique used to map interactions between distant DNA elements (Lieberman-Aiden et al., 2009, Rao et al., 2014). Its more recent derivative, Capture Hi-C (CHi-C), allows for high-depth characterisation of DNA interactions in loci of interest (Dryden et al., 2014). CHi-C has been applied to gene promoters in multiple blood cell types (Javierre et al., 2016) and to GWAS loci in diseases such as cancer (Baxter et al., 2018, Dryden et al., 2014, Jager et al., 2015) and autoimmune conditions (Martin et al., 2015). HiChIP also builds on Hi-C by enriching for interactions that colocalise with an immunoprecipitated chromatin fraction (such as that marked by histone 3 lysine 27 acetylation, a hallmark of active chromatin) (Mumbach et al., 2016). HiChIP was recently applied in primary T cells (Mumbach et al., 2017) and B cells, in the context of systemic lupus erythematosus (SLE) (Pelikan et al., 2018). To empirically determine the function of GWAS SNPs, direct perturbation is now widely carried out using the CRISPR-Cas9 system, either through genome editing or chromatin activation/interference (CRISPRa/CRISPRi) of promoters and enhancers (Adli, 2018).

Here, we apply these novel technologies in the context of psoriasis to identify gene targets and subsequently annotate a psoriasis risk locus at 9q31 where the closest gene, Krüppel-like factor 4 (*KLF4*), encodes a transcription factor with a range of relevant functions including skin barrier formation (Segre et al., 1999) and immune signalling (An et al., 2011), but is situated more than 500 kb from the lead SNP for the GWAS signal (Tsoi et al., 2012).

## Results

### Capture Hi-C identified novel gene targets in psoriasis susceptibility loci

We generated sequencing data for three region CHi-C experiments in HaCaT and My-La cells, in biological duplicate. In HaCaT we generated CHi-C data for both unstimulated and IFN-γ-stimulated cells (Supplementary Table 2). Our overarching design targeted genetic regions associated with several immune-mediated diseases including psoriasis (see Methods). We aimed for 10,000 and obtained an average of 11,508 mapped Hi-C fragments (di-tags) per bait fragment with a mean capture efficiency of 70%. Capture Hi-C Analysis of Genomic Organisation (CHiCAGO) was used to identify significant interactions for each cell type within the unique di-tags. For the My-La cell line we noted a large number of significant *trans* interactions (CHiCAGO score ≥ 5) spanning different chromosomes (7,329/55,700 total interactions from all captured immune-mediated disease loci). We found that the majority of these (around 71%) mapped to interactions between two known translocated loci in My-La cells (Netchiporouk et al., 2017). In light of this, the interactions were filtered to only include cis (same-chromosome) interactions.

We filtered the CHi-C interactions to include only those involving psoriasis GWAS loci; we had targeted 104 lead GWAS SNPs at genome-wide significance and their associated proxy SNPs at r^2^ > 0.8, corresponding to 907 HindIII bait fragments (Supplementary Table 1). Across the three capture experiments, we obtained an average of 7,075 interactions (CHiCAGO score ≥ 5) originating from targeted psoriasis fragments (Supplementary Table 2). The data were enriched for long-range interactions, with more than 70% of the significant interactions in the psoriasis loci spanning >100 kb. The median interaction distances were comparable between cell types: 227 kb (HaCaT unstimulated), 234 kb (HaCaT stimulated) and 297 kb (My-La) (Supplementary Figure 1).

To validate our CHi-C data we overlaid the interactions with a published expression quantitative trail locus (eQTL) dataset, in which the lead psoriasis SNP had been colocalised with the lead eQTL SNP in CD4+ T cells and monocytes (Raj et al., 2014). We hypothesised that long-distance eQTL-gene promoter pairings would often implicate chromatin looping. The study reported 15 lead GWAS SNPs with 26 corresponding lead eQTL proxy SNPs, of which 16 proxies, representing 9 lead GWAS SNPs, overlapped baited fragments in our study (the lack of complete overlap mostly owed to our study prioritising index SNPs from more recent meta-analyses over older studies; in addition, we were not able to design baits for one of the reported regions at rs7552167; see Methods and Supplementary Table 4). Eight of the proxies were captured within a HindIII fragment that contained, or was within 20 kb of, the eQTL gene itself (Supplementary Table 3). A further seven proxies were within, or adjacent to, fragments that showed evidence of interacting with the distal eQTL gene in our cell line CHi-C data (CHiCAGO score ≥ 5) (Supplementary Table 3). Only one distal proxy, rs8060857, did not show any evidence of interacting with the eQTL gene (*ZNF750*); this was the most distant eQTL gene at approximately 720 kb away (Supplementary Table 3). Therefore, this is strong evidence that our CHi-C data can show links between distal functional GWAS SNPs and their target gene, even across different cell types.

In all the cell lines, approximately 30% of the interactions occurred between the psoriasis bait fragment and a fragment containing a transcription start site (Ensembl 75) (Supplementary Table 4). The total number of interacting gene targets was 442 in unstimulated HaCaT cells, 461 in stimulated HaCaT cells and 761 in My-La cells, comprising a set of 934 genes. Of these, 295 gene targets (31.6%) were shared between all cell types, whilst 50, 63 and 386 targets were unique in unstimulated HaCaT, IFN-γ-stimulated HaCaT and My-La cells, respectively. Unstimulated and stimulated HaCaT cells shared a large proportion of their gene targets (355 targets; 77-80%). Bait fragments that interacted with genes tended to interact with multiple promoter-containing fragments corresponding to different genes; a median of 2 fragments in HaCaT cells (unstimulated or stimulated) and 3 fragments in My-La cells, implicating between 2-4 genes (Supplementary Figure 2). This demonstrated a complex relationship between implicated enhancers and interacting genes, consistent with previously-reported findings (Martin et al., 2015, Mifsud et al., 2015, Mumbach et al., 2017)

Of all the genes interacting with psoriasis-associated loci, 580 (62% of the total set of 934 genes) were classified as protein-coding by Ensembl. We used RNA-seq to determine gene expression and found that approximately 46% of genes that were long-range targets of a CHi-C psoriasis-implicated bait were also expressed in the same cell type. These included compelling candidates such as *IL23A*, *PTGER4*, *STAT3* and *NFKBIZ* (Supplementary Table 5). In addition, 129 genes (48% of all genes overlapping baited fragments) that overlapped baited fragments were expressed in the cell lines (Supplementary Table 5).

Stimulating HaCaT cells with IFN-γ caused the differential expression of 535 genes (adjusted P < 0.10): 88 down-regulated and 447 up-regulated (Supplementary Table 6). Whilst the down-regulated genes were not enriched for any biological pathways, the up-regulated genes were enriched for 196 biological processes that included such psoriasis-relevant GO terms as “GO:0045087 innate immune response” (P = 9.39 x 10^-20^), “GO:0034097 response to cytokine stimulus” (P = 7.32 x 10^-15^) and “GO:0034340 response to type I interferon” (P = 1.08 x 10^-10^) (Supplementary Table 7). Twelve of the differentially expressed genes overlapped a psoriasis capture bait fragment (Supplementary Table 8) and included *ERAP1, ERAP2, IFIH1, RNF114, SOCS1* and *STAT2*. In addition, 12 differentially-expressed genes were involved in bait-promoter interactions (Supplementary Table 8) and included candidates such as *ICAM1*, *KLF4* and *STAT3*. However, the vast majority of these differentially-expressed genes interacted similarly with the psoriasis-associated baits in both unstimulated and stimulated cells (CHiCAGO score ≥ 5) (Supplementary Table 8).

### Examples of CHi-C interactions implicating target genes for psoriasis

We found that the CHi-C data represent three broad scenarios across psoriasis loci: (i) the interactions lend evidence to support the nearest or reported gene candidate; (ii) the interactions suggest distal or novel gene candidates; or (iii) the interactions add complexity to the locus, highlighting multiple gene candidates. Here, we present some examples to represent each scenario.

#### CHi-C implicates the nearest or reported gene candidate

At the intergenic locus 9q31, the psoriasis association falls between two distant gene clusters where the suggested gene candidate was Krüppel-like factor 4 (*KLF4*) due to its relevant biological functions in differentiation and innate immunity (Tsoi et al., 2012). Our CHi-C data showed significant interactions (CHiCAGO score ≥ 5) between the psoriasis association and the promoter of *KLF4* in both unstimulated and stimulated HaCaT cells, over a distance of approximately 560 kb (Figure 1A). In both unstimulated and stimulated HaCaT cells, one bait fragment (chr9:110810592-110816598) interacted with *KLF4*. In stimulated HaCaT cells, a second bait fragment (chr9:110798319-110798738) also interacted, coinciding with a more than five-fold increase in *KLF4* expression (FC = 5.78; adj. P = 4.26 x 10^-8^). In My-La cells a similar conformation was observed; however, the interactions did not coincide with the fragment containing the gene itself. Furthermore, *KLF4* expression was not detected by RNA-seq in My-La cells, suggesting a cell-type specific mechanism. In all cell types, long-range interactions also stretched from the psoriasis locus to the telomeric side of the gene desert but fell short of the nearest gene on that side (*ACTL7B*) by approximately 35 kb.

**Figure 1.**
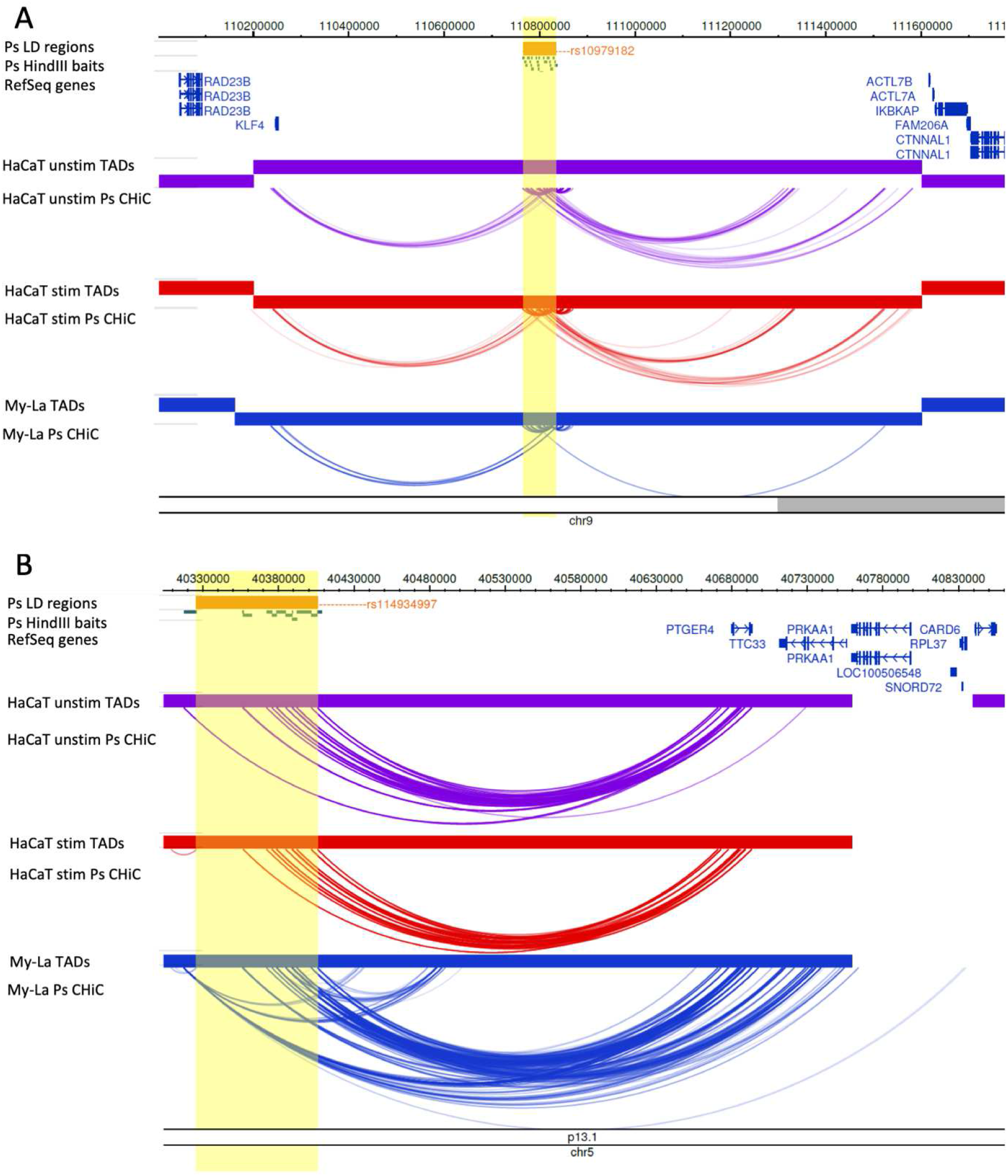
Examples of CHi-C Interactions implicating nearest/reported genes. Interactions are shown in the 9q31.2 (*KLF4*) locus (A) and the 5p13.1 (*PTGER4*) locus (B). The tracks include psoriasis (Ps) LD blocks as defined by SNPs in r^2^ >0.8 with the index SNP, baited HindIII fragments, RefSeq genes (NCBI), TADs (shown as bars) and CHi-C interactions significant at CHiCAGO score ≥ 5 (shown as arcs) in three conditions: unstimulated HaCat cells (purple), HaCaT cells stimulated with IFN-γ (red) and My-La cells (blue). The highlighted region indicates the psoriasis LD block in each locus. The figure was made with the WashU Epigenome Browser, GRCh37/hg19 (Zhou et al., 2013).

At the 5p13.1 locus, the psoriasis SNPs are similarly intergenic (Tsoi et al., 2015) but the nearest gene *PTGER4* has been shown to be a strong candidate for other autoimmune diseases at this locus (Mumbach et al., 2017). Our CHi-C data showed interactions (CHiCAGO score ≥ 5) between multiple psoriasis-associated fragments and *PTGER4* over approximately 300 kb to the other end of the TAD; a finding that was robust in all cell types (Figure 1B). *PTGER4* expression was detected by RNA-seq in all cell types (Supplementary Table 5). In My-La cells, interactions also stretched to the promoters of *TTC33*, which was expressed in all cell types, and *RPL37*, for which expression was not detected in any cell type.

#### CHi-C implicates distal or novel gene candidates

At the 2p15 locus, the psoriasis association tagged by rs10865331 was originally assigned to the nearest gene *B3GNT2*; however, the CHi-C interactions skipped *B3GNT2* (~120 kb downstream) and instead implicated the promoter of Copper Metabolism Domain Containing 1 (*COMMD1*), a gene involved in NFkB signalling, over approximately 435 kb upstream (Figure 2A) (Maine et al., 2007, Tsoi et al., 2012). This interaction occurred in stimulated HaCaT cells and My-La cells, and *COMMD1* expression was detected by RNA-seq in all cell types (Supplementary Table 5). *B3GNT2* expression was also detected in all cell types (data not shown).

**Figure 2.**
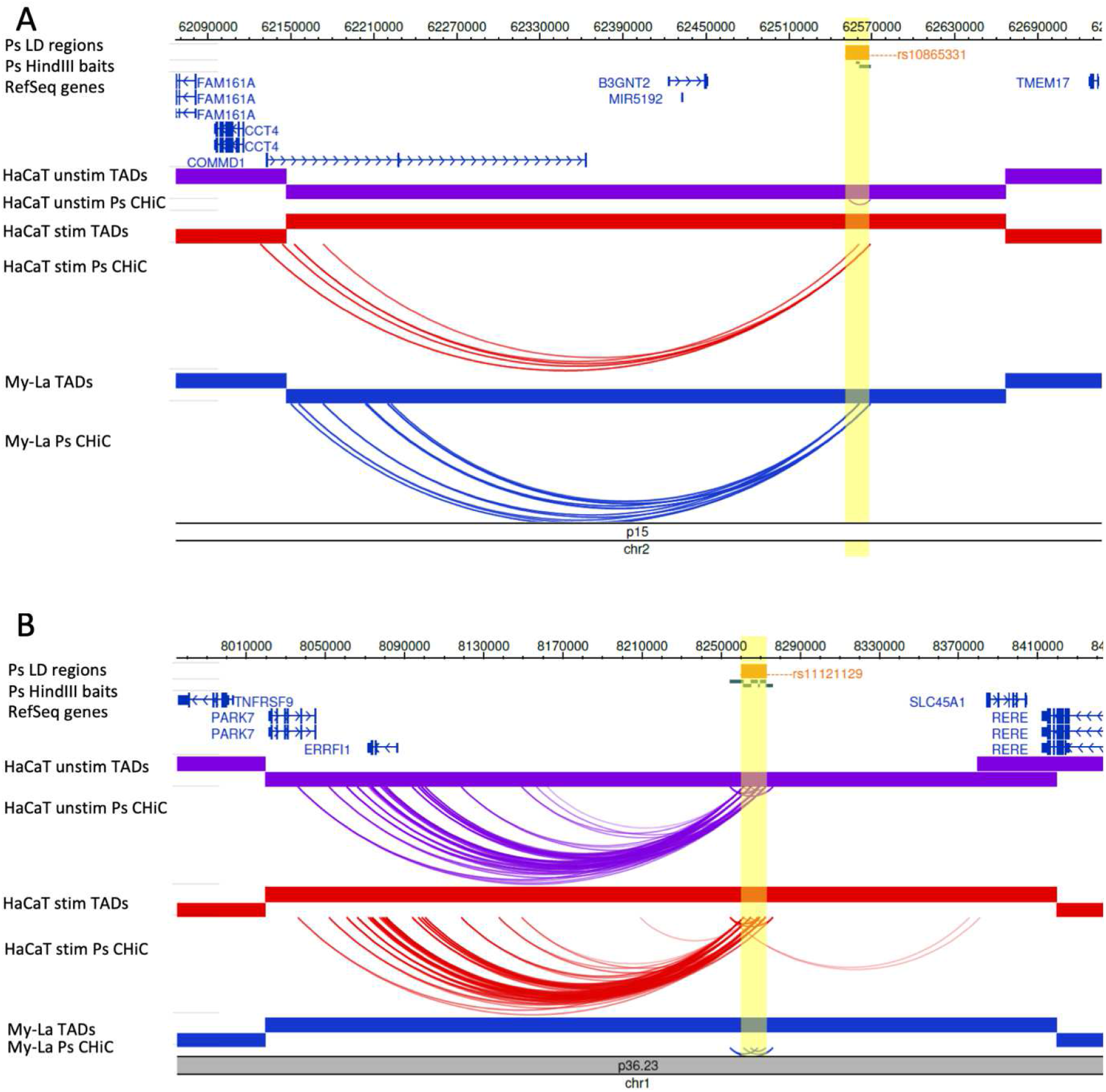
Examples of CHi-C interactions in gene deserts implicating distal/novel genes. Interactions are shown in the 2p15 (*B3GNT2*) locus (A) and the 1p36.23 (*RERE*, *SLC45A1*, *ERRFI1*, *TNFRSF9*) locus (B). The tracks include psoriasis (Ps) LD blocks as defined by SNPs in r^2^ >0.8 with the index SNP, baited HindIII fragments, RefSeq genes (NCBI), TADs (shown as bars) and CHi-C interactions significant at CHiCAGO score ≥ 5 (shown as arcs) in three conditions: unstimulated HaCat cells (purple), HaCaT cells stimulated with IFN-g (red) and My-La cells (blue). The highlighted region indicates the psoriasis LD block in each locus. The figure was made with the WashU Epigenome Browser, GRCh37/hg19 (Zhou et al., 2013).

At the 1p36.23 locus, the association tagged by rs11121129 is closest to *SLC45A1*, and was originally assigned to multiple putative gene targets (Tsoi et al., 2012). However, the CHi-C data showed interactions (CHiCAGO score ≥ 5) between the psoriasis LD block and the promoter of ERBB Receptor Feedback Inhibitor 1 (*ERRFI1*), an important regulator of keratinocyte proliferation and differentiation, in both unstimulated and stimulated HaCaT cells (Figure 2B). This interaction was not observed in My-La cells and moreover, *ERRFI1* expression was detected in HaCaT cells (unstimulated and stimulated) but not My-La cells. An interaction between the psoriasis association and the promoter of *SLC45A1* was also observed in stimulated, but not unstimulated, HaCaT cells (Figure 2B), however *SLC45A1* expression was not detected by RNA-seq in any of the cell lines.

#### CHi-C adds complexity to the locus

At the 6p22.3 locus, the psoriasis signal tagged by rs4712528 is intronic to *CDKAL1*, and there were 11 psoriasis-associated intronic fragments that also interacted with the *CDKAL1* promoter in My-La cells (Figure 3A); *CDKAL1* expression was detected in all cells (Stuart et al., 2015). However, there were also long-range interactions (CHiCAGO score ≥ 5) between psoriasis-associated fragments and *SOX4* over 950 kb in all cell types (Figure 3A). *SOX4* is a compelling gene candidate with roles in IL17A production and skin inflammation in mice (Malhotra et al., 2013); here *SOX4* expression was detected in HaCaT cells but not in My-La cells.

**Figure 3.**
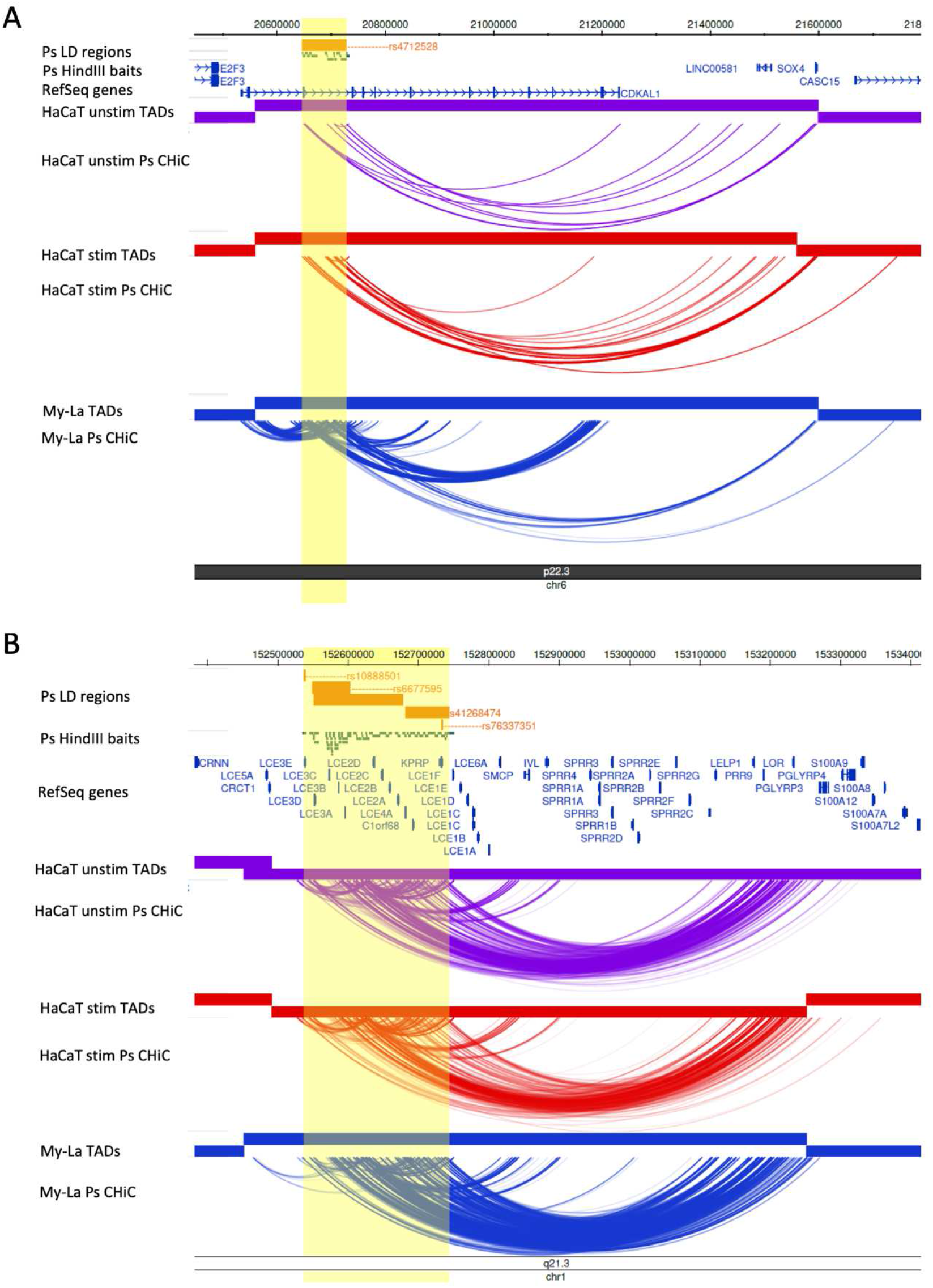
Examples of CHi-C interactions adding complexity to a locus. Interactions are show in the 6p22.3 (*CDKAL1*) locus (A) and the 1q21.3 (*LCE3B*, *LCE3C*) locus (B). The tracks include psoriasis (Ps) LD blocks as defined by SNPs in r^2^ >0.8 with the index SNP, baited HindIII fragments, RefSeq genes (NCBI), TADs (shown as bars) and CHi-C interactions significant at CHiCAGO score ≥ 5 (shown as arcs) in three conditions: unstimulated HaCat cells (purple), HaCaT cells stimulated with IFN-g (red) and My-La cells (blue). The highlighted region indicates the psoriasis LD block in each locus. The figure was made with the WashU Epigenome Browser, GRCh37/hg19 (Zhou et al., 2013).

At the 1q21.3 locus, multiple risk SNPs are located at the late cornfield envelope (LCE) gene cluster in the epidermal differentiation complex (EDC). One of the associations in this locus is a 32 kb deletion that removes the *LCE3B* and *LCE3C* genes (de Cid et al., 2009, Li et al., 2011, Tsoi et al., 2012). The CHi-C data showed multiple, robust interactions between the psoriasis-associated regions at the LCE genes, including from within the 32kb LCE3C/B-del region, and genes downstream in the EDC that included *IVL*, *LOR*, *PRR9* and *SPRR* genes, over a distance of ~600 kb (Figure 3B). Of these genes, *IVL* interacted with psoriasis baits in unstimulated but not stimulated HaCaT cells and its expression decreased upon stimulation (FC = 0.40; adj P = 0.0139). The coding genes directly interacting with fragments within the 32 kb deletion were *LCE3A*, *PRR9*, *LELP1*, *SPRR2B* and *SPRR2C*. Of these, only expression of proline-rich region 9 (*PRR9*) was detected; in HaCaT cells but not in My-La cells. *PRR9* was previously shown to be upregulated in psoriatic plaques and induced by IL17A and so may be an important distal gene target in this locus (Swindell et al., 2016).

### The 9q31 psoriasis risk locus forms long-range interactions with KLF4 and harbours likely-regulatory variants

We focused our attention on the large intergenic locus at 9q31, tagged by rs10979182, which showed long-range interactions between the psoriasis-associated SNPs and *KLF4* (Figure 1A) (Tsoi et al., 2012). *KLF4* expression was also upregulated by IFN-γ (Supplementary Table 6), suggesting that it may be an important player within an inflammatory environment. We wanted to determine if any functional enhancer-promoter relationship existed between the SNPs and *KLF4* or other, distal, genes in the locus.

First, we characterised the psoriasis-associated SNPs in 9q31 by mining publicly available epigenetic datasets and tools. There are ninety variants in tight LD (r^2^ > 0.8) with the lead GWAS SNP rs10979182 (1KG Phase 3 European) (Figure 4A); several of which intersect modified histone marks (H3K4me1 and H3K27ac) in several cell types from ENCODE, corresponding with four putative enhancer elements overlapping H3K4me1 and H3K27ac occupancy (Figure 4B). In primary human keratinocyte (NHEK) cells, enhancer histone marks were most prominent in enhancers 2-4 (Figure 4C). The SNPs also overlap DNase hypersensitivity sites and transcription factor binding sites (Figure 4C), that correspond with enhancer elements in NHEK according to ChromHMM (Ernst and Kellis, 2012).

**Figure 4:**
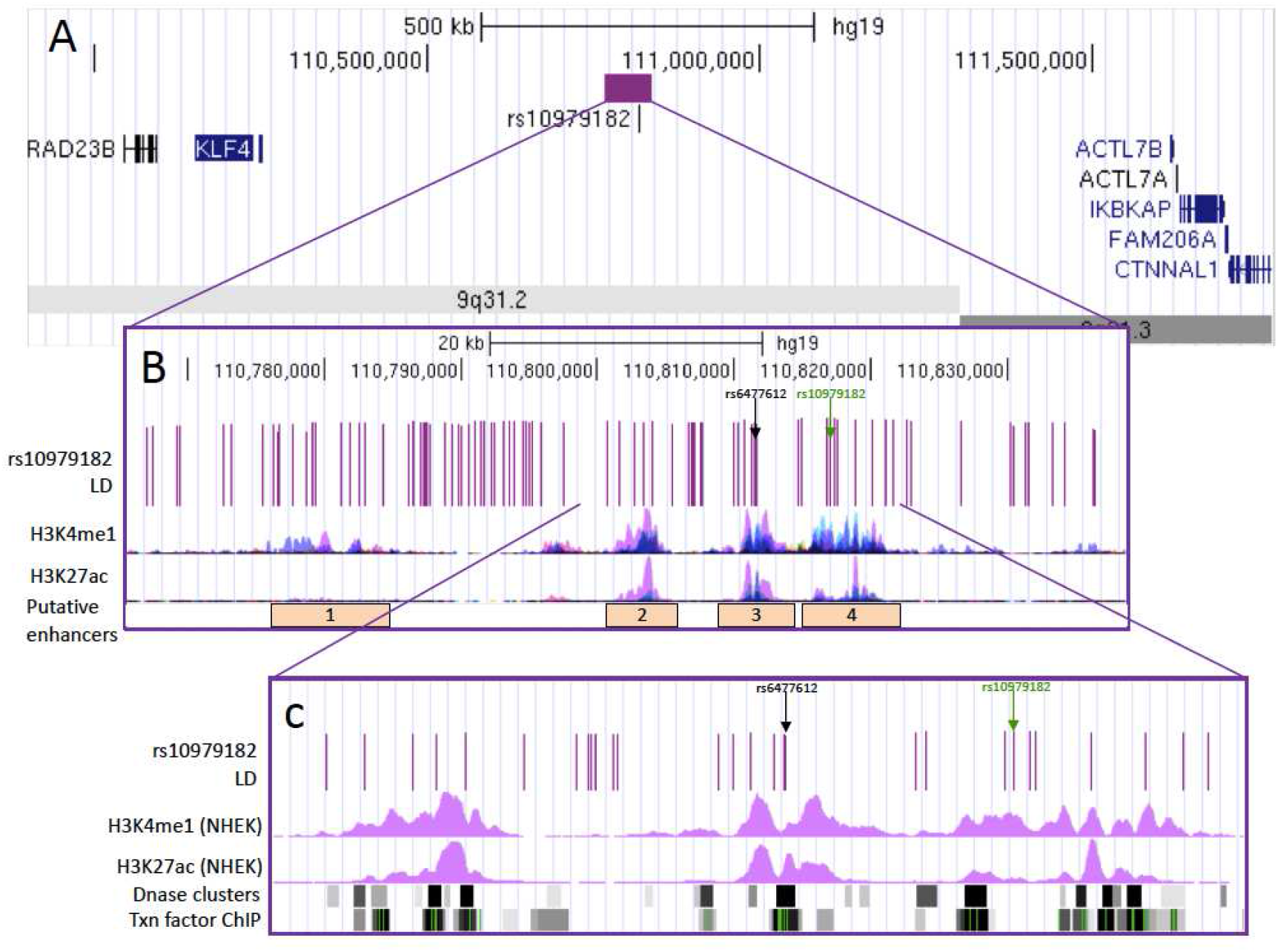
Overview of SNPs in LD with rs10979182 overlaying four putative enhancer elements in the 9q31.2 locus. (A) The purple bar demonstrates the location of the rs10979182 LD block (r^2^ > 0.8) in the ~1Mb gene desert between two gene clusters, shown by UCSC genes (Hsu et al., 2006). (B) The 90 SNPs in LD with rs10979182 are denoted by purple lines and H3K4me1 and H3K27ac ChIP-seq tracks from ENCODE are shown as peaks in GM12878 (red), H1-hESC (yellow), HSMM (green), HUVEC (light blue), K562 (dark blue), NHEK (purple) and NHLF (pink) cells (Dunham et al., 2012). The index SNP, rs10979182, is shown as a green arrow and the most likely regulatory SNP, rs6477612, is shown as a black arrow. (C) Zoom-in of the putative enhancers 2-4 showing SNPs overlaying ENCODE regulatory marks: H3K4me1 and H3K27ac ChIP-seq (NHEK), DNase clusters, and transcription factor ChIP clusters across 91 cell types as grey/black bars, where darkness indicates signal strength. For ChIP clusters, green lines indicate the highest scoring site of a Factorbook-identified canonical motif for the corresponding factor. The index SNP, rs10979182, is shown as a green arrow and the most likely regulatory SNP, rs6477612, is shown as a black arrow. The figure was made with the UCSC Genome Browser, GRCh37/hg19 (Kuhn et al., 2013).

No eQTLs were identified in the set according to Haploreg v4.1. RegulomeDB identified rs6477612, situated within the third putative enhancer, as the SNP with the highest putative regulatory potential with a score of 2a. rs6477612 is in tight LD (r^2^ = 0.92, 1KG EUR) with rs10979182 and was located within the HindIII fragment found to interact with *KLF4* in HaCaT cells in our CHi-C data (chr9:110810592-110816598; hg19), making it a prioritised SNP of interest.

### HiChIP data suggested that the interactions between KLF4 and psoriasis SNPs are active in HaCaT cells, but not My-La cells

As a complementary approach to CHi-C, we used the recently developed HiChIP method to identify H3K27ac-mediated interactions in our cell lines. We examined the significant HiChIP interactions contacting the promoter of *KLF4*. In HaCaT cells, there were H3K27ac peaks at the *KLF4* promoter and in abundance across the gene desert (Figure 5). In contrast, there were a lack of H3K27ac peaks in My-La cells in 9q31 and, correspondingly, no significant HiChIP interactions. This lack of H3K27ac occupancy indicates a differential activation state in this region between HaCaT and My-La cells. In HaCaT cells, the H3K27ac peak at *KLF4* interacted with multiple H3K27ac peaks across the gene desert, including peaks at psoriasis-associated enhancers 3 and 4 and at the previously-published interacting region from the breast cancer study in both unstimulated and stimulated HaCaT cells (Figure 5) (Dryden et al., 2014).

**Figure 5.**
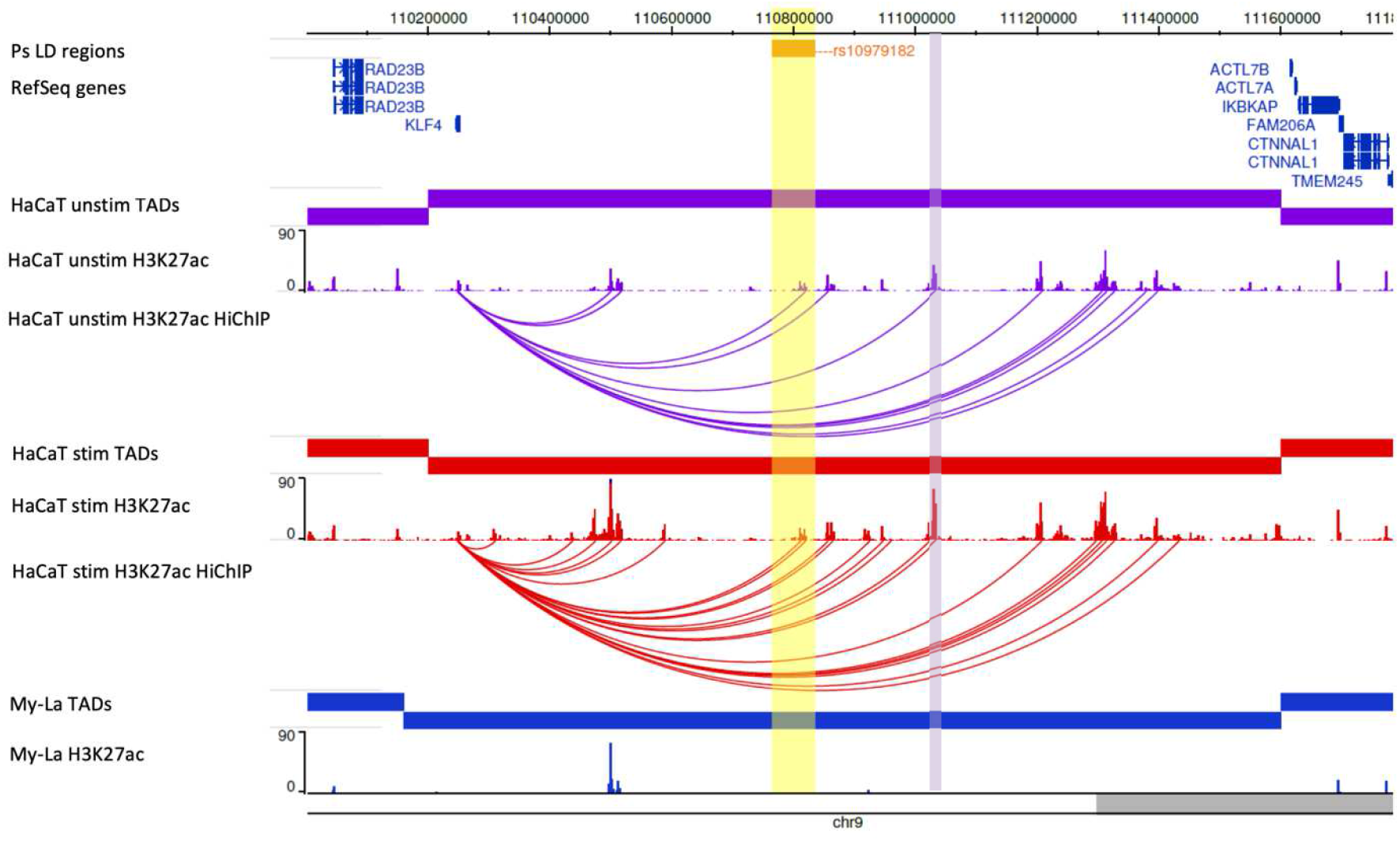
HiChIP (H3K27ac) interactions with the *KLF4* promoter in the 9q31 locus. The tracks include: psoriasis LD block as defined by SNPs in r^2^ >0.8 with rs10979182, RefSeq genes, TADs (shown as bars), H3K27ac occupancy (shown as peaks), and significant HiChIP interactions (shown as arcs) in three conditions: unstimulated HaCaT cells (purple), HaCaT cells stimulated with IFN-g (red) and My-La cells (blue). The HiChIP interactions were restricted to those originating from the *KLF4* promoter and filtered to include those with at least 5 reads. The yellow highlighted region indicates the psoriasis LD block at rs10979182. The purple highlighted region indicates the previously described *KLF4*-interacting region in the breast cancer study (Dryden et al., 2014). The figure was made with the WashU Epigenome Browser, GRCh37/hg19 (Zhou et al., 2013).

Restricting the significant H3K27ac-mediated interactions to those originating from the psoriasis-associated enhancers in HaCaT cells revealed multiple enhancer-enhancer interactions in 9q31. However, other than *KLF4* there were no other gene targets implicated by H3K27ac-mediated HiChIP interactions in 9q31 (data not shown).

We observed an increase in the number and strength of H3K27ac peaks in the 9q31 TAD (chr9:110,202,281-111,602,280) in stimulated HaCaT cells compared with unstimulated HaCaT cells (Figure 5). The number of peaks increased from 60 to 77, and there was a significant increase in the median peak signal from ~5.3 to ~9.5 in shared peaks (P < 0.0001, Wilcoxon matched-pairs signed rank test). Consequently, there were double the number of H3K27ac-mediated interactions with the *KLF4* promoter in stimulated cells (Figure 5). This also corresponded with an over 5-fold upregulation of gene expression upon IFN-γ stimulation in HaCaT cells (FC = 5.78; adj. P = 4.26 x 10^-8^). Combined, this suggests that there is more enhancer activity in 9q31 in stimulated cells than unstimulated cells, and correlates with promoter CHi-C data from other groups suggesting that there is a strong relationship between gene expression and the absolute number of enhancer interactions (Burren et al., 2017).

### 3C-qPCR supplemented HiChIP/CHi-C findings in the 9q31 locus

We used 3C-qPCR in an effort to confirm the interaction between the psoriasis-associated putative enhancer 3 (rs6477612) and *KLF4* to further prioritise regulatory SNPs. Our 3C experiment utilised both the enhancer and the *KLF4* gene as focus anchors, in both HaCaT and My-La cell lines. The enhancer-focused 3C experiment identified interaction peaks with regions approximately 2.5kb and 8.7kb downstream of *KLF4* in My-La, and with the downstream 8.7kb region alone in HaCaT (Supplementary Figure 3).

The *KLF4*-focused 3C experiment showed that *KLF4* significantly interacted with several intergenic psoriasis-associated fragments, including the fragment containing the third putative enhancer (rs6477612), in HaCaT cells, but not in My-La cells (Supplementary Figure 4). This corroborates the CHi-C data, which showed a more robust interaction between the enhancer and the *KLF4* gene in HaCaT cells. A positive control interaction linking a distal breast cancer-associated locus with *KLF4* (Dryden et al., 2014, Baxter et al., 2018) demonstrated the strongest interaction with the *KLF4* promoter region in both cell types (Supplementary Figure 4).

Taken together, the 3C results confirm a close spatial proximity between the psoriasis-associated SNPs and *KLF4* in 9q31. However, there is no clear peak of interaction among the LD block that would implicate some SNPs over others. In addition, stronger interactions were seen between *KLF4* and regions further upstream in the gene desert, which correlates with previous CHi-C findings in breast cancer cells (Dryden et al., 2014) and Hi-C findings in NHEK cells (Rao et al., 2014) (illustrated in Supplementary Figure 5).

### ChIP-qPCR confirmed binding of regulatory histone marks in 9q31 in HaCaT cells

We performed ChIP-qPCR of the histone marks H3K4me1 and H3K27ac to determine cell-type specificity of enhancer activity within the *KLF4*-interacting psoriasis loci. Primers were designed to target 150-200 bp regions encompassing predicted peaks of H3K27ac occupancy in the four putative enhancers identified from ENCODE data (NHEK). H3K4me1 and H3K27ac occupancy was detected at all tested loci in HaCaT and My-La cells (Figure 6A). However, occupancy was significantly increased in HaCaT cells with an enrichment of both H3K4me1 and H3K27ac at enhancer 3 (H3K4me1 P = 0.0372, H3K27ac P < 0.0001) and H3K27ac at enhancer 4 (P < 0.0001) in HaCaT cells in comparison with My-La cells (Figure 6A). Stimulation of HaCaT cells with IFN-γ had little effect on the occupancy of H3K4me1 or H3K27ac at the regions tested within the enhancers, or at a region tested at the *KLF4* promoter (Figure 6B).

**Fig. 6.**
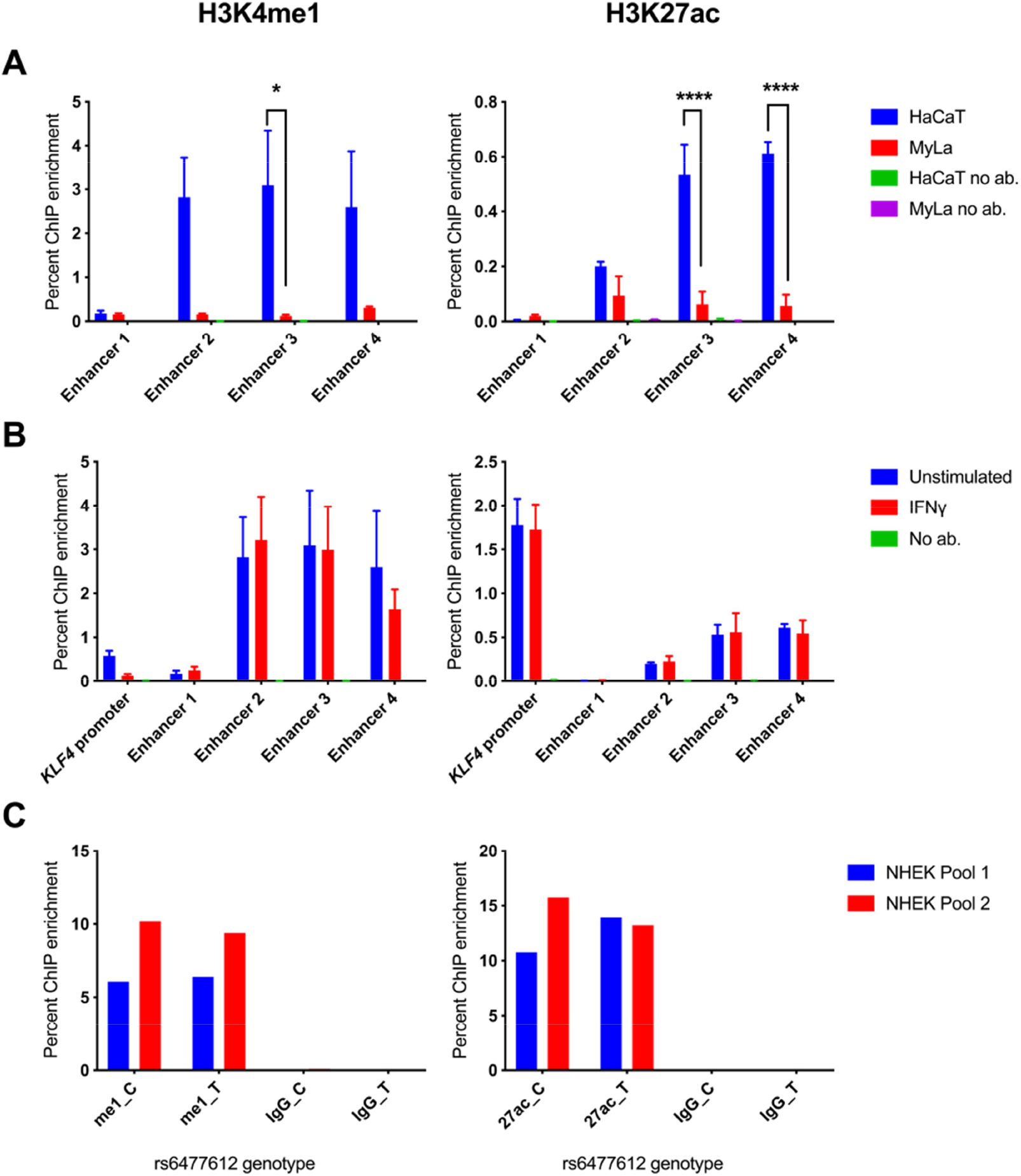
ChIP-qPCR for modified histone marks H3K4me1 and H3K27ac in 9q31. A) Enhancer peaks defined by H3K27ac binding in ENCODE NHEK data were targeted in HaCaT cells (blue columns) and My-La cells (red columns). B) Enhancer peaks were targeted in unstimulated (blue) and stimulated (red) HaCaT cells. Graphs show mean ChIP enrichment of triplicate ChIP libraries ± SEM, and samples with no antibody are consistently included for comparison, although they are often too low to be visible. To identify differential ChIP enrichment, 2-way ANOVA tests were performed in GraphPad prism using Sidak’s multiple comparisons test. Asterisks denote adjusted P < 0.05. C) Allele-specific ChIP-qPCR for H3K27ac and H3K4me1 at rs6477612 in NHEK cells. Chromatin from two separate pools of NHEK cells, each comprising cells from three individual donors, was immunoprecipitated with H3K27ac (27ac), H3K4me1 (me1) or non-specific IgG antibody (IgG) and qPCR was conducted using a TaqMan genotyping assay for rs6477612 detecting C (risk) or T (protective) alleles. Percentage ChIP enrichment was calculated by comparing the signal for each allele in the immunoprecipitated DNA with the signal for each allele in the input DNA for each of the two samples.

To determine potential effects of the risk or protective allele of rs6477612, the most likely regulatory SNP within enhancer 3, we performed allele-specific ChIP at rs6477612 for H3K4me1 and H3K27ac in two pools of NHEK cells. However, there was no discernible difference in H3K4me1 or H3K27ac occupancy at the risk (C) or protective (T) allele of rs6477612 (**Figure 6C**).

In summary, by combining the HiChIP, CHi-C 3C and ChIP evidence we could determine that the psoriasis-associated enhancer region interacts with *KLF4* in both My-La and HaCaT but is only active in HaCaT cells. Enhancer activity in 9q31 is increased after IFN-g stimulation, correlating with an increase in *KLF4* gene expression, although we were unable to detect increases in H3K27ac occupancy at the tested psoriasis-associated enhancer regions.

### CRISPR activation suggested that the psoriasis-associated enhancer elements regulate KLF4 expression in 9q31

We employed CRISPRa in 9q31 in order to determine whether activating the psoriasis-associated enhancers could impact on gene expression (*KLF4* or other, distal genes), implicating a functional role for the long-range interactions. Pools of single guide RNA (sgRNA) targeting SNPs within the four psoriasis-associated enhancers were introduced into HaCaT cells stably expressing the CRISPR activator dCas9-P300 (see Supplementary Methods, Figure 1 for overview of sgRNA locations). Across the CRISPRa experiments we found that activating the enhancers caused a significant increase in *KLF4* expression of approximately 2-fold in comparison with the control, scrambled sgRNA (enhancer 1 P= 0.0005, enhancer 2 P= 0.0196, enhancer 3 P = 0.0001 and enhancer 4 P=0.0001 respectively) (Figure 7). Pool 3, targeting enhancer 3 containing the most likely regulatory SNP rs6477612, had the greatest impact with a 2.2-fold increase in *KLF4* expression. In addition, IkappaB Kinase Complex-Associated Protein (*IKBKAP*) to the telomeric end of the gene desert was also subtly but significantly upregulated by approximately 1.2-fold in cell lines containing sgRNA pools 1, 3 and 4, in comparison with the scrambled sgRNA (P= 0.0088, P=0.0239 and P= 0.0188, respectively; one-way ANOVA) (Figure 7). We found that *FAM206A* and *CTNNAL1* were not significantly affected by CRISPRa. The remaining two genes *ACTL7A* and *ACTL7B*, were not detectable in any HaCaT cell line, transduced or otherwise (data not shown).

**Figure 7.**
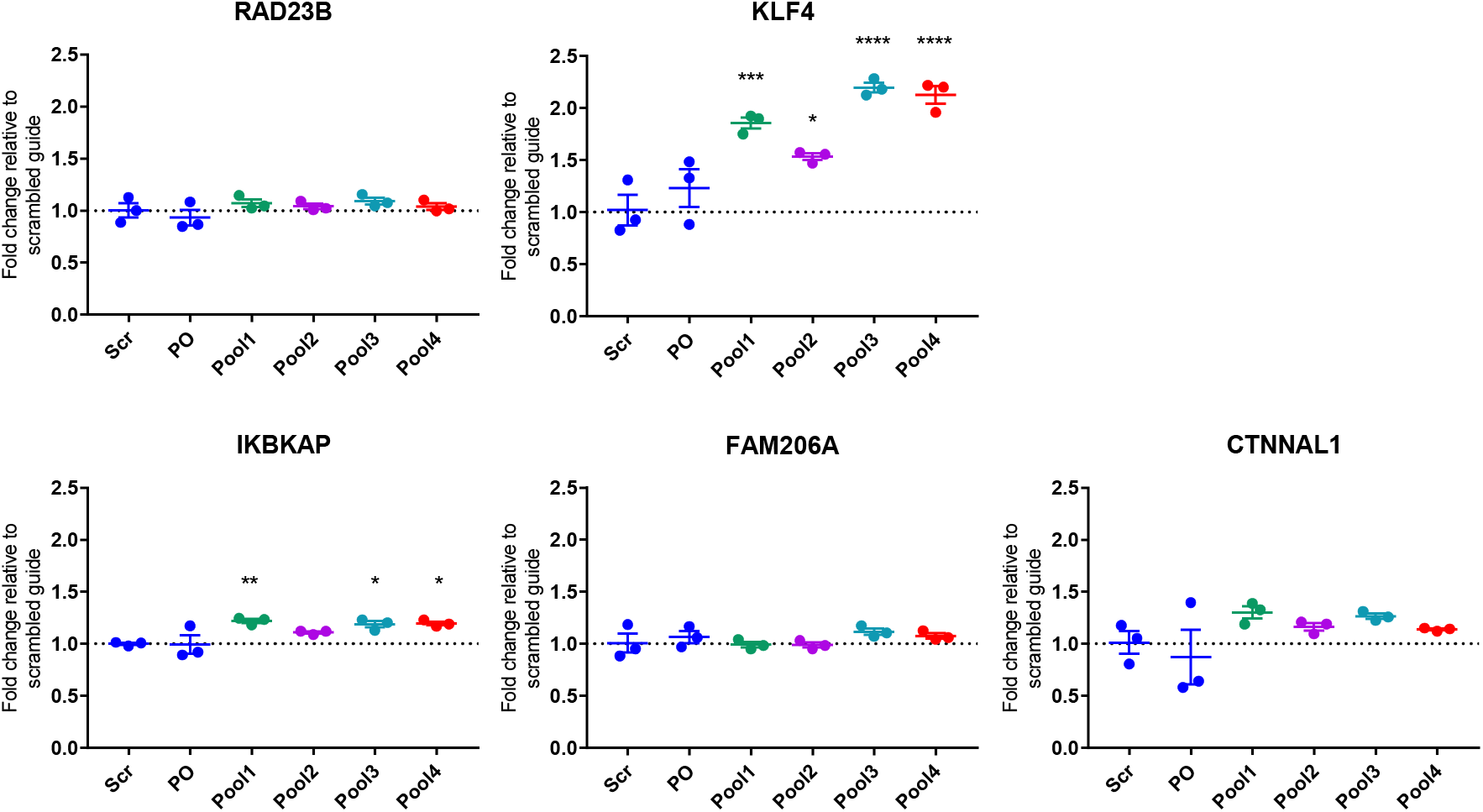
qPCR results for genes within the 9q31 locus in HaCaT cells expressing dCas9-P300. HaCaT cells expressing dCas9-P300 were transduced with pools of plasmids containing sgRNA targeting psoriasis SNPs (pools 1-4), a scrambled sgRNA (Scr) or the same plasmid without a specific guide cloned in (plasmid only; PO). TaqMan qPCR results are shown for *RAD23B*, *KLF4*, *IKBKAP*, *FAM206A* and *CTNNAL1*. Housekeeping genes used were *TBP* and *YWHAZ*. For statistical analysis, a one-way ANOVA was performed for each gene, comparing each condition with the scrambled guide, using Dunnett’s multiple comparisons test. Asterisks denote P < 0.05. Graphs show the mean fold-change in comparison with the scrambled guide, ±SEM of biological triplicate cell lines.

To determine the transcriptome-wide effects of activating the psoriasis-associated enhancers in 9q31, RNA-seq was performed on the HaCaT dCas9-P300 cells expressing the sgRNA in Pool 3 (putative enhancer 3) and compared with cells expressing the scrambled sgRNA (Supplementary Table 9). In line with the qPCR experiment, RNA-seq revealed an approximately 3-fold increased expression of *KLF4* in the Pool 3 cells. (FC = 2.92; adj P = 0.0546). There was an approximately 1.3-fold increase in *IKBKAP* expression, but this was not significant (FC = 1.26; adj P = 0.7331).

The RNA-seq analysis showed that there were an additional 236 differentially expressed genes in the CRISPRa experiment (adjusted P ≤ 0.10); 128 up-regulated and 108 down-regulated (Supplementary Table 9). Importantly, CRISPRa of the psoriasis-implicated enhancer in this keratinocyte cell line not only resulted in an increase in *KLF4* expression, but a differential expression of 3 keratin genes (4, 13 and 15), confirming the importance of this enhancer and the *KLF4* gene in skin cell function. Keratin 4 was the most differentially expressed gene in our data with Keratin 15 being the 6^th^ most differentially expressed. Previous studies also demonstrated a differential impact of *KLF4* on keratin gene regulation (Chen et al., 2003, Okano et al., 2000, He et al., 2015).

Confirming the importance of *KLF4* in skin cells, and validating previous findings of differential gene expression with *KLF4* stimulation, differential genes also included *EREG* (an epidermal growth factor) *MMP13* (extracellular matrix protein gene) and *CLDN8* (claudin 8; important in epithelium tight junctions). We also demonstrated a ~10-fold reduction in expression of *ALPG*, from a family of alkaline phosphatases showing the largest fold change in a previous *KLF4* over-expression study (Chen et al., 2003) (Supplementary Table 9).

The up-regulated genes were enriched in several biological pathways according to the GAGE analysis, of which the most significant related to RNA processing (Supplementary Table 10). According to the STRING database, differential genes were enriched in a number of relevant pathways including apoptosis and response to cell stress, emphasising the role that *KLF4* plays in cell cycle regulation and supporting previous findings demonstrating that over-expressing *KLF4* leads to G1/S cell cycle arrest (Chen et al., 2001) (Supplementary Table 11).

## Discussion

Genetic predisposition is the largest known risk factor for psoriasis. The GWAS era has provided a wealth of genetic loci associated with psoriasis, and yet if we are to exploit these data fully, for clinical benefit with new and more effective therapies, there is a requirement for a better understanding of their biological significance. Recent functional studies have used sophisticated post-GWAS technologies to assign gene targets, cell types and functional mechanisms in loci associated with other, related complex conditions (Martin et al., 2015, McGovern et al., 2016, Wang et al., 2013, Simeonov et al., 2017). Here, we have combined these technologies for the first time focused on investigation into the functional genomics of psoriasis.

Our study provides findings that are complementary to previously published data. For example, our data in cell lines demonstrates a long-range interaction between psoriasis-associated SNPs and *PTGER4* in the intergenic psoriasis risk locus at 5p13.3, similar to published, promoter CHi-C data in several other primary cell types, including psoriasis-relevant cells such as macrophages, monocytes, CD4+ T cells, CD8+ T cells and neutrophils (Javierre et al., 2016). Recently, HiChIP again demonstrated how this locus forms functional enhancer interactions with *PTGER4* (Mumbach et al., 2017). We also demonstrate how chromatin confirmation data can provide evidence for suspected causal gene targets or provide support for regions that show an eQTL to a putative causal gene.

We provide a compelling pathway to integrate data from public resources, combined with chromatin conformation, molecular biology and genome editing techniques, to link disease associated variants to causal genes, cell types, mechanism and pathways. This is illustrated with the intergenic 9q31 psoriasis risk locus. Here we implicate *KLF4* as the likely causal psoriasis risk gene in this region, by showing how the gene is a cell type-specific target for regulation by disease associated variants. Crucially, by incorporating RNA-seq analysis, we demonstrate that activating the disease implicated enhancers also activates downstream pathways of *KLF4*, showing how perturbation of a regulatory sequence and modest effect on transcription factor expression can have a profound effect on downstream gene targets in disease. Our findings also complement a study showing that the *KLF4* transcription factor is dysregulated in psoriatic skin (Kim et al., 2014). KLF4 protein contains both an activation and repression domain and is known to either upregulate or down regulate pathways in a tissue and context-dependent manner; therefore, its role in disease is likely to be complex (Ray, 2016).

The CRISPRa experiment showed that the psoriasis-associated enhancers predominantly affected *KLF4* expression, but there was a small but significant effect also on the expression of *IKBKAP*; a gene previously implicated in NFκB signalling (Cohen et al., 1998). This is perhaps not surprising, since the CHi-C data suggested that the chromatin conformation in 9q31 would bring the psoriasis-associated enhancers into close proximity with the genes to the telomeric side of the gene desert. The RNA-seq experiment confirmed an upregulation of *IKBKAP*, indicating that these enhancers, and maybe other GWAS implicated regulatory regions, may have multiple downstream effects.

In conclusion, we provide evidence as to gene targets in psoriasis risk loci, supporting assigned candidates and in some regions suggesting novel candidates. We also focus on a specific risk locus and, by moving from associated variants to gene, cell type mechanism and pathway, we demonstrate how *KLF4* is a likely gene target of the GWAS association in 9q31. This investigative pathway is applicable to all GWAS studies and may be the next pivotal step towards patient benefit and clinical translation.

## Methods

### Cell culture

The spontaneously transformed keratinocyte cell line HaCaT (Addexbio) was cultured in high-glucose Dulbecco’s modified eagle’s medium (DMEM) supplemented with 10% foetal bovine serum (FBS) and 1% penicillin-streptomycin (Thermo Fisher Scientific, final concentration 100 U penicillin, 0.1 mg streptomycin/mL). For stimulation experiments, the media was supplemented with 100 ng/mL recombinant human IFN-γ (285-IF-100; R&D Systems) and cells were incubated for 8 hours prior to harvest. Pools of adult Normal Human Epidermal Keratinocytes (NHEK; PromoCell) were cultured in Keratinocyte Growth Medium 2 (PromoCell) supplemented with 0.06 mM CaCl2. For chromatin-based experiments, HaCaT and NHEK cells were crosslinked for 10 minutes in 1% formaldehyde and the cross-linking reaction was quenched with 0.135M glycine.

The cancer-derived human CD8+ T-lymphocyte cell line My-La CD8+ (Sigma-Aldrich) was cultured in Roswell Park Memorial Institute (RPMI) 1640 medium supplemented with 10% AB human serum (Sigma Aldrich), 100 U/mL recombinant human IL-2 (Sigma-Aldrich) and 1% penicillin-streptomycin (final concentration 100 U penicillin, 0.1 mg streptomycin/ml). For chromatin-based experiments, My-La cells were crosslinked for 10 minutes in 1% (ChIP, HiChIP) or 2% (CHi-C, 3C) formaldehyde. Crosslinking reactions were quenched with 0.135M glycine.

Lenti-X 293T cells (Takara Biosciences) were used for lentivirus production. They were cultured in DMEM high-glucose supplemented with 10% FBS and penicillin streptomycin (final concentration 100 U penicillin, 0.1 mg streptomycin/ml).

### Capture Hi-C

For CHi-C, RNA baits were designed to target all known non-MHC psoriasis risk loci, defined by one or more independent SNPs associated with psoriasis in GWAS (Supplementary Table 1). The total number of SNPs included was 107 (59 associated with Europeans, 42 with Chinese, and 6 associated with both European and Chinese cohorts) corresponding with 68 loci. The baits were selected to target HindIII fragments that overlapped with the linkage disequilibrium (LD) block in each locus, defined by SNPs in r^2^ > 0.8 with the lead SNP (1000 Genomes Phase 3 release, European). Due to sequence restraints, baits could not be designed for the 1p36.11 (rs7552167, rs4649203) (Cheng et al., 2014, Tsoi et al., 2012) and 1q31.1 (rs10789285) (Tsoi et al., 2015) loci; therefore in total there were 104 SNPs corresponding with 66 psoriasis risk loci in the final capture library (907 HindIII fragments).

The psoriasis baits were combined with a capture library targeting multiple GWAS loci across several immune-mediated diseases: juvenile idiopathic arthritis, asthma, psoriatic arthritis, rheumatoid arthritis and systemic sclerosis. The majority of these baits were included in our previous region CHi-C experiment (Martin et al., 2015); results from these loci are not described in the present study. Each 120 bp bait was targeted to within 400 bp of a HindIII fragment end, comprised 25-65% GC content and contained fewer than three unknown bases. The baits were synthesised by Agilent Technologies.

CHi-C libraries were generated in biological duplicate for My-La and HaCaT (unstimulated or stimulated) cells according to previously described protocols (Dryden et al., 2014, Martin et al., 2015); see Supplementary Methods. Briefly, 50 million crosslinked cells were lysed and the chromatin digested with HindIII, followed by biotinylation and in-nucleus ligation. Crosslinks were reversed, the DNA was purified and biotin-streptavidin pulldown was used to enrich for ligation sites. The libraries were amplified and capture was performed using the RNA baits described above using the SureSelect reagents and protocol (Agilent). Following a second amplification the libraries were sequenced using paired-end Illumina SBS technology (see Supplementary Methods for details).

CHi-C sequence data were processed through the Hi-C User Pipeline (HiCUP) v0.5.8 (Wingett et al., 2015). For each cell type, the two biological replicates were simultaneously run through CHiCAGO v1.1.8 (Cairns et al., 2016) in R v3.3.0 and significant interactions were called with a score threshold of 5. Subsequently, the significant interactions were restricted to those between loci on the same chromosome (cis). BEDTools v2.17.0 (Quinlan, 2014) was used to detect interactions between psoriasis-associated fragments and gene promoters, defined by fragments covering regions within 500 bp of transcription start sites (Ensembl release 75; GRCh37). CHi-C interactions were visualised using the WashU Epigenome Browser (Zhou et al., 2013).

### Hi-C

For each cell type, a pre-CHi-C library was generated (see Supplementary Methods). Hi-C libraries were sequenced using paired-end Illumina SBS technology (see Supplementary Methods for details). The sequence data was filtered and adapters removed using fastp v0.19.4 (Chen et al., 2018). The reads were then mapped to the GRCh38 genome with Hi-C Pro v2.11.0 (Servant et al., 2015), using default settings. The Hi-C interaction matrices were normalised within Hi-C Pro using iterative correction and eigenvector decomposition (ICE). Topologically associating domains (TADs) were then called in TADtool software (Kruse et al., 2016) using insulation score with the normalised Hi-C contact matrices, binned at a 40 kb resolution. TADs were visualised alongside CHi-C interactions on the WashU Epigenome Browser (Zhou et al., 2013).

### HiChIP

HiChIP libraries were generated according to the Chang Lab protocol (Mumbach et al., 2016); see Supplementary Methods. Briefly, 10 million crosslinked cells were lysed, digested with MboI, biotinylated and ligated. Immunoprecipitation was performed with H3K27ac antibody, after which the DNA was purified and de-crosslinked followed by biotin-streptavidin pulldown. Tagmentation was performed using Tn5 (Illumina) and the libraries were amplified with Nextera indexing primers (Illumina). Sequencing was performed using paired-end Illumina SBS technology (see Supplementary Methods for details).

Sequencing data for the HiChIP libraries was filtered and the adapters were removed using fastp v0.19.4. The reads were then mapped to the GRCh38 genome with Hi-C Pro v2.11.0, using default settings. Long-range interactions were called using hichipper v0.7.3 (Lareau and Aryee, 2018). The anchors were called using the self-circle and dangling end reads corresponding to each sample. The remainder of the settings were left default. Interactions were filtered by FDR < 0.10 and a minimum of 5 reads. The interactions were filtered to those originating from the H3K27ac peak on the *KLF4* promoter before being uploaded for visualisation on the WashU Epigenome Browser (Zhou et al., 2013). H3K27ac ChIP-seq tracks were generated using the self-circle and dangling end reads from the libraries and extended for 147 bp. The datasets were scaled according to read depth before being uploaded for visualisation to the WashU Epigenome Browser (Zhou et al., 2013).

To compare H3K27ac signal in shared peaks between unstimulated and stimulated HaCaT cells in 9q31, genome-wide anchors reported from hichipper were first combined to produce a merged peak set. Then the signal from the two conditions was intersected on the peaks using BEDTools map function and the mean signal for each peak was reported for each condition. The resulting values were imported in R and normalized using DESeq2 estimate size factors function (Love et al., 2014). The normalized counts for peaks within the 9q31 TAD (chr9:110,202,281-111,602,280) were compared between the two conditions using a Wilcoxon matched-pairs signed rank test in GraphPad Prism.

### RNA-seq

3’ mRNA sequencing libraries were generated for cell lines using the QuantSeq 3’ mRNA-Seq Library Prep Kit FWD for Illumina (Lexogen). RNA-seq libraries were generated for unstimulated HaCaT cells (N = 4), stimulated HaCat cells (N = 3) and My-La cells (N = 1). Libraries were sequenced using single-end Illumina SBS technology. Reads were quality trimmed using Trimmomatic v0.38 (Bolger et al., 2014) using a sliding window of 5 with a mean minimum quality of 20. Adapters and poly A/poly G tails were removed using Cutadapt v1.18 (Martin, 2011) and then UMIs were extracted from the 5’ of the reads using UMI-tools v0.5.5 (Smith et al., 2017). Reads were then mapped using STAR v2.5.3a (Dobin et al., 2013) on the GRCh38 genome with GENCODE annotation v29. Reads were de-duplicated using UMIs with UMI-tools and then counted using HTSeq v0.11.2 (Anders et al., 2015). Count matrixes were analysed in R 3.5.1 and normalisation and differential expression analysis was conducted using DESeq2 v1.22.2. Differentially expressed genes were called with an adjusted P value of 0.10 (FDR 10%). Gene set enrichment Pathway analysis was performed using GAGE v2.32.1 (Luo et al., 2009) using “normal” shrinked log fold changes from DESeq2. For detection of expressed genes in the cell lines, we considered RNA-seq counts greater than 1 in at least one of the sequenced samples.

### Functional annotation in 9q31

SNPs in LD (r^2^>0.8) with the lead SNP rs10979182 were examined for their intersection with ENCODE data for histone marks, transcription factor binding sites and DNase hypersensitivity. RegulomeDB v1.1 (Boyle et al., 2012) was used to rank the SNPs based on likely regulatory function. The SNPs were also assessed using Haploreg v4.1 (Ward and Kellis, 2012), which includes expression quantitative trait loci (eQTL) data from several studies including GTEx (Lonsdale et al., 2013) and GEUVADIS (Lappalainen et al., 2013).

### 3C-qPCR in 9q31

3C libraries were generated in biological triplicate as previously described (Naumova et al., 2012). Briefly, 20-30 million crosslinked cells were lysed, digested, ligated and purified as in the initial Hi-C steps, omitting biotinylation. Control libraries were constructed using bacterial artificial chromosome (BAC) clones as described by (Naumova et al., 2012). Eleven minimally-overlapping BAC sequences were selected to span the 9q31 locus (chr9:110168556-111889073, hg19); see Supplementary Methods.

qPCR was carried out using SYBR Green or TaqMan technology to determine interaction frequencies in the 9q31 locus. Relative interaction frequencies were calculated using the BAC curve and normalised to a short-range control. Significant interactions (P < 0.05) were detected using one-way ANOVA in GraphPad Prism with Dunnett’s or Tukey’s test for multiple comparisons.

### ChIP-qPCR in 9q31

Chromatin immunoprecipitation (ChIP) libraries were generated as previously described (McGovern et al., 2016); see Supplementary Methods. Briefly, 10 million cross-linked cells were lysed and the chromatin was fragmented and immunoprecipitated with antibodies for H3K4me1 (Abcam ab8895) or H3K27ac (Abcam ab4729). ChIP enrichment was measured at loci of interest by qPCR using SYBR Green or TaqMan technology. The data were normalised by calculating the percentage of total chromatin that was immunoprecipitated in comparison with an input sample.

For SYBR experiments, primers were designed using Primer3 (http://primer3.ut.ee/) (Untergasser et al., 2012) to target regions of 100-200 bp encompassing enhancers in 9q31 (Supplementary Methods). A TaqMan SNP genotyping probe (Applied Biosystems, assay ID C__29343482_10) was used for an allele-specific analysis at rs6477612, detecting C (risk) or T (protective). The difference in antibody binding to each allele was determined by the percentage of ChIP enrichment in comparison with the signal for each allele obtained from the input sample. Differences in ChIP enrichment were calculated in GraphPad Prism using two-way ANOVA (P < 0.05). Experiments were performed in biological triplicate (HaCaT and My-La lines) or duplicate (NHEK cells).

### CRISPR activation in 9q31

CRISPR activation using the dCas9-P300 complex was performed in HaCaT cells to determine the role of the four putative enhancers in 9q31. To select sgRNA in 9q31, SNPs in r^2^> 0.8 with rs10979182 were prioritised by their overlap with enhancer elements, defined by active regulatory regions in NHEK according to ChromHMM (Ernst et al., 2011). sgRNA sequences were designed using the online CRISPOR tool (Concordet and Haeussler, 2018) to target loci within 200 bp of the prioritised SNPs (mean = 85 bp). sgRNA were selected based on specificity score and proximity to the SNP. In total there were 27 SNPs; two of these could not be targeted by sgRNA within 200 bp (rs7019552 and rs11355519) and another two SNPs, rs4979624 and rs7029094, were captured by a single sgRNA targeting the intervening region. In total, therefore, there were 24 sgRNA; these were grouped into four pools of 5-7 SNPs to target the four putative enhancers (see Supplementary Methods).

A HaCaT cell line stably expressing dCas9-P300 was generated by lentiviral transduction with the CRISPRa plasmid pLV_dCas9-p300-P2A-PuroR plasmid (Addgene 83889) (Klann et al., 2017). Next, the sgRNA sequences were cloned into the pLKO5.sgRNA.EFS.GFP plasmid (Addgene 57822) (Heckl et al., 2014) and equimolar plasmid pools were generated for each enhancer. The plasmid pools were then packaged using the same lentiviral method as the dCas9-P300 plasmid (Supplementary Methods). The guide pools were introduced into the stable HaCaT dCas9-P300 cells using a second round of lentiviral transduction and cells that had integrated the sgRNA plasmids were isolated by flow cytometry for GFP. To assay gene expression, qPCR was performed on extracted RNA using the TaqMan RNA-to-Ct 1-step kit (Thermofisher Scientific) with TaqMan assays for the following genes in the 9q31 locus: *RAD23B*, *KLF4*, *ACTL7A*, *ACTL7B*, *IKBKAP*, *FAM206A* and *CTNNAL1* (assay IDs detailed in Supplementary Methods). Delta-delta Ct analysis was conducted against a control HaCaT dCas9-P300 cell line transduced with the sgRNA plasmid containing a previously-published scrambled, non-targeting insert (Scramble2; (Lawhorn et al., 2014)). Two housekeeping gene assays, *TBP* and *YWHAZ*, were used for normalisation (Supplementary Methods). For the CRISPRa Pool with the greatest impact on *KLF4* expression, RNA-seq and gene set enrichment analyses were performed as described above. In addition, differentially expressed genes were processed through the STRING database to identify potential protein-protein interaction networks (Szklarczyk et al., 2015).

## Supporting information

supplementary results

## Declarations

## Acknowledgements

The authors would like to acknowledge the assistance given by IT Services and the use of the Computational Shared Facility at The University of Manchester.

## Data availability

The sequence datasets generated and analysed during the current study are available in the GEO repository under accession number GSE137906.

## Authors’ contributions

All authors contributed to the preparation of the manuscript. SE, RBW, KD and HRJ contributed to the conception and design of the experiment, HRJ, KD, AMG, PM, CS, JH, OG, CT, JD, VPG, YF and PG to the acquisition or analysis of the data, HRJ, SE, RBW, KD, CS, AY, APM, AA, PM and GO to the interpretation of the results.

