## Supplementary methods for "Chromatin-based techniques map DNA interaction landscapes in psoriasis susceptibility loci and highlight *KLF4* as a target gene in 9q31"

### Cell culture

HaCaT keratinocyte cells were obtained from Addexbio (T0020001); these are *in vitro* spontaneously transformed keratinocytes from histologically normal skin. Cells were cultured in high-glucose Dulbecco’s modified eagle’s medium (DMEM) supplemented with 10% foetal bovine serum (FBS) and penicillin-streptomycin (Thermo Fisher Scientific, final concentration 100 U penicillin, 0.1 mg streptomycin/ml). For HaCaT stimulation experiments, the media was supplemented with 100 ng/mL recombinant human IFN-γ (285-IF-100; R&D Systems) and cells incubated for 8 hours prior to harvest.

Pools of adult Normal Human Epidermal Keratinocytes (NHEK) were obtained from PromoCell (C-12006) and cultured in Keratinocyte Growth Medium 2 (PromoCell) supplemented with 0.06 mM CaCl_2_.

My-La CD8+ cells were obtained from Sigma-Aldrich (95051033). These cells are cancerous human T-lymphocytes derived from a patient with mycosis fungoides. Cells were cultured in Roswell Park Memorial Institute (RPMI) 1640 medium supplemented with 10% AB human serum (Sigma Aldrich), 100 U/mL recombinant human IL-2 (Sigma-Aldrich) and penicillin-streptomycin (final concentration 100 U penicillin, 0.1 mg streptomycin/ml)..

Lenti-X 293T cells were obtained from Takara Biosciences (632180). These cells are a sub-clone of the human embryonic kidney (HEK) cell line and are optimised for viral protein production. 293T cells were cultured in DMEM high-glucose supplemented with 10% FBS and penicillin streptomycin (final concentration 100 U penicillin, 0.1 mg streptomycin/ml).

### Cell crosslinking for chromatin-based experiments

HaCaT and NHEK cells were crosslinked for 10 minutes in 1% formaldehyde and the reaction was quenched with 0.135M glycine. The crosslinked cells were pelleted, washed in PBS and the supernatant removed. Cells were snap frozen on dry ice and stored at -80°C. My-La cells were crosslinked for 10 minutes in 1% (ChIP, HiChIP) or 2% (Hi-C, 3C) formaldehyde. The reaction was quenched with 0.135M glycine, the supernatant removed and the cells snap frozen on dry ice and stored at -80°C.

### Capture Hi-C

For CHi-C, RNA baits were designed to target all known non-MHC psoriasis risk loci, defined by one or more independent SNPs associated with psoriasis in GWAS (Supplementary Table 1). The total number of SNPs included was 107 (59 associated with Europeans, 42 with Chinese, and 6 associated with both European and Chinese cohorts) corresponding with 68 loci. The baits were selected to target HindIII fragments that overlapped with linkage disequilibrium (LD) block in each locus, defined by SNPs in r^2^ > 0.8 with the lead SNP (1000 Genomes Phase 3 release, European). Baits could not be designed for the 1p36.11 (rs7552167, rs4649203) (Cheng et al., 2014; L. C. Tsoi et al., 2012) and 1q31.1 (rs10789285) (Lam C. Tsoi et al., 2015) loci; therefore there were 104 SNPs corresponding with 66 psoriasis loci in the final capture library (907 HindIII fragments).

The psoriasis baits were combined with a capture library targeting multiple GWAS loci across several immune-mediated diseases: juvenile idiopathic arthritis, asthma, psoriatic arthritis, rheumatoid arthritis and systemic sclerosis. The majority of these baits were included in our previous region capture Hi-C experiment (P. Martin et al., 2015). This initial capture library was applied to My-La Hi-C libraries, whereas a slightly updated capture library, that included some additional baits, was applied to HaCaT Hi-C libraries. The two different baitmaps included the same psoriasis regions, and results from non-psoriasis loci are not described in the present study. In addition, a control locus, which represents a well-characterised region of long-range interactions, *HBA*, was also included (174.57 kb genomic; 26 restriction fragments; 6.71 kb/restriction fragment). Each 120 bp bait was targeted to within 400 bp of a HindIII fragment end, comprised 25-65% GC content and contained fewer than three unknown bases. The baits were synthesised by Agilent Technologies.

CHi-C libraries were generated in biological duplicate for My-La and HaCaT (unstimulated or stimulated) cells according to previously described protocols (Dryden et al., 2014; P. Martin et al., 2015). 50 million crosslinked cells were lysed and the chromatin digested with HindIII at 37°C overnight. Restriction cut sites were filled in using dCTP, dGTP, dTTP and biotin-14-dATP (Life Technologies), then in-nucleus ligation was carried out at 16°C for 4-6 hours. Crosslinks were removed by proteinase-K overnight at 65°C and RNA was digested using RNaseA for 60 minutes at 37°C. The DNA was purified by sequential phenol and phenol-chloroform extractions and ethanol-precipitated at -20°C overnight, followed by two further phenol-chloroform extractions and a second overnight precipitation.

A 40 µg aliquot of DNA was taken forward for further processing following QC steps. T4 DNA polymerase used to remove biotin-14-dATP from non-ligated ends then the DNA purified by phenol-chloroform extraction and ethanol precipitation overnight. The DNA was sheared using a Covaris S220 sonicator and end-repair was performed using T4 DNA polymerase, T4 DNA polynucleotide kinase and DNA polymerase I, large (Klenow) fragment. The sample was purified using Qiagen MinElute Kit, with a modified protocol described by (Belton et al., 2012). Klenow (exo-) was used to adenylate DNA fragment ends and a double-sided SPRI bead size selection was used to obtain fragments of approximately 200-600 bp. Dynabeads MyOne Streptavidin C1 beads (Life Technologies) were used to pull down biotinylated fragments, which were then ligated to annealed Illumina sequencing adapters. PCR was performed using Phusion HF (NEB) and TruPE PCR primers (Illumina), then the amplified DNA was cleaned twice using 1.8X volume of SPRI beads.

Amplified DNA up to 750 ng was concentrated using a vacuum concentrator and bound to the capture baits in a single hybridisation reaction using SureSelectXT reagents and protocol by Agilent Technologies. The biotinylated baits were captured using Dynabeads MyOne Streptavidin T1 beads (Life Technologies). Following washes, the libraries were amplified on the beads using Phusion HF and barcoded TruPE primers then the amplified DNA cleaned twice using 1.8X volume of SPRI beads. The quality and quantity of the capture Hi-C libraries was tested by Bioanalyzer and KAPA qPCR (Kapa Biosystems). Capture Hi-C libraries were analysed by 75 bp paired-end Next Generation Sequencing on an Illumina NextSeq500 at the Genomic Technologies Core Facility at the University of Manchester (My-La) or HiSeq 4000 at Edinburgh Genomics at the University of Edinburgh (HaCaT).

CHi-C sequence data were processed through the Hi-C User Pipeline (HiCUP) v0.5.8 (Wingett et al., 2015). For each cell type, the two biological replicates were simultaneously run through Capture Hi-C Analysis of Genomic Organisation (CHiCAGO) v1.1.8 (Cairns et al., 2016) in R v3.3.0 and significant interactions were called with a score threshold of 5. Subsequently, the significant interactions were restricted to those between loci on the same chromosome (cis). BEDTools v2.17.0 (Quinlan, 2014) was used to detect interactions between psoriasis-associated fragments and gene promoters, defined by fragments covering regions within 500 bp of transcription start sites (Ensembl release 75; GRCh37). CHi-C interactions were visualised using the WashU Epigenome Browser (Zhou et al., 2013).

## Hi-C

For each cell type, a single Hi-C library was generated by re-amplifying the pre-capture Hi-C library bound to streptavidin beads, using Phusion HF and barcoded TruPE primers. The amplified DNA was cleaned twice using 1.8X volume of SPRI beads. The quality and quantity of the Hi-C libraries was tested by Bioanalyzer and KAPA qPCR. Hi-C libraries were analysed by Next Generation Sequencing. The My-La Hi-C library was sequenced on an Illumina HiSeq 2500 generating 100bp paired ends at the Babraham Institute Sequencing Facility, Cambridge. The HaCaT Hi-C libraries were sequenced on an Illumina HiSeq 4000 generating 75 bp paired ends at Edinburgh Genomics at the University of Edinburgh. The sequence data was filtered and the adapters were removed using fastp v0.19.4 (Chen, Zhou, Chen, & Gu, 2018). The reads were then mapped to the GRCh38 genome with Hi-C Pro v2.11.0 (Servant et al., 2015), using default settings. The Hi-C interaction matrices were normalised within Hi-C Pro using iterative correction and eigenvector decomposition (ICE). Topologically associating domains (TADs) were then called in TADtool software (Kruse, Hug, Hernández-Rodríguez, & Vaquerizas, 2016) using insulation score with the normalised Hi-C contact matrices, binned with 40 kb resolution. TADs were visualised alongside CHi-C interactions on the WashU Epigenome Browser (Zhou et al., 2013).

### HiChIP

HiChIP libraries were generated according to the Chang Lab protocol (Mumbach et al., 2016). Briefly, 10 million crosslinked cells were lysed and the chromatin digested using 375 U of MboI (NEB, R0147M) for 4 hours at 37°C. Fragment ends were filled in using dCTP, dGTP, dTTP and biotin-14 dATP (Life Technologies) and ligated at room temperature overnight. The nuclei were lysed and the chromatin sheared to lengths of approximately 200-700 bp using a Covaris S220. Immunoprecipitation was performed overnight at 4°C using 20 µg of H3K27ac antibody (Abcam ab4729). The DNA was captured on a 1:1 mixture of protein A and G Dynabeads (Invitrogen 10001D and 10003D). After washes, the DNA was eluted with proteinase K at 65°C overnight. The sample was cleaned using Zymo Clean and Concentrator Columns (Zymo D4013) and quantified using the Qubit DNA HS kit. 20-35 ng of DNA was taken forward for biotin-pulldown with streptavidin C-1 beads at room temperature for 30 minutes. The beads were suspended in TD buffer from the Nextera kit and transposed with Tn5 (Illumina) at 55°C for exactly 10 minutes. The volume of Tn5 was dependent on DNA quantity and defined by the original HiChIP protocol (Mumbach et al., 2016). After washes, the library was amplified off the beads using Phusion polymerase and Nextera indexing primers (Illumina). SPRI beads were used to select fragments approximately 300 – 700 bp in length. Quantification and quality control of the final HiChIP library was conducted using a Bioanalyzer and KAPA quantification kit (Kapa Biosystems). Libraries underwent Next Generation Sequencing on a HiSeq 2500 at The Babraham Institute Sequencing facility, Cambridge, generating 100 bp paired-ends.

Sequencing data for the HiChIP libraries was filtered and the adapters were removed using fastp v0.19.4. The reads were then mapped to the GRCh38 genome with Hi-C Pro v2.11.0, using default settings. Long-range interactions were called using hichipper v0.7.3 (Lareau & Aryee, 2018). The anchors were called using only self-circle and dangling end reads corresponding to each sample. The remainder of the settings were left default. Interactions were filtered by FDR < 0.10 and a minimum of 5 reads. The interactions were filtered to those originating from the H3K27ac peak on the *KLF4* promoter before being uploaded for visualisation on the WashU Epigenome Browser (Zhou et al., 2013). H3K27ac ChIP-seq tracks were generated using only self-circle and dangling end reads from the libraries and extended for 147 bp. The datasets were scaled according to read depth before being uploaded for visualisation to the WashU Epigenome Browser (Zhou et al., 2013).

To compare H3K27ac signal in shared peaks between unstimulated and stimulated HaCaT cells in 9q31, genome-wide anchors reported from hichipper were first combined to produce a merged peak set. Then the signal from the two conditions was intersected on the peaks using BEDTools map function and the mean signal for each peak was reported for each condition. The resulting values were imported in R and normalized using DESeq2 estimate size factors function (Love et al., 2014). The normalized counts for peaks within the 9q31 TAD (chr9:110,202,281-111,602,280) were compared between the two conditions using a Wilcoxon matched-pairs signed rank test in GraphPad Prism.

### RNA-seq

3’ mRNA sequencing libraries were generated for cell lines using the Lexogen QuantSeq 3’ mRNA-Seq Library Prep Kit FWD for Illumina. RNA-seq libraries were generated for unstimulated HaCaT cells (N = 4), stimulated HaCat cells (N = 3) and My-La cells (N = 1). Libraries were sequenced using single-end Illumina SBS technology. Reads were quality trimmed using Trimmomatic v0.38 (Bolger, Lohse, & Usadel, 2014) using a sliding window of 5 with a mean minimum quality of 20. Adapters and poly A/poly G tails were removed using Cutadapt v1.18 (M. Martin, 2011) and then UMIs were extracted from the 5’ of the reads using UMI-tools v0.5.5 (Smith, Heger, & Sudbery, 2017). Reads were then mapped using STAR v2.5.3a (Dobin et al., 2013) on the GRCh38 genome with GENCODE annotation v29. Reads were de-duplicated using UMIs with UMI-tools and then counted using HTSeq v0.11.2 (Anders, Pyl, & Huber, 2015). Count matrixes were analysed in R 3.5.1 and normalisation and differential expression analysis was conducted using DESeq2 v1.22.2 (Love, Huber, & Anders, 2014). Differentially expressed genes were called with an adjusted P value of 0.10 (FDR 10%). Gene set enrichment Pathway analysis was performed using GAGE v2.32.1 (Luo, Friedman, Shedden, Hankenson, & Woolf, 2009) using “normal” shrinked log fold changes from DESeq2. For detection of expressed genes in the cell lines, we considered RNA-seq counts greater than 1 in at least one of the sequenced samples.

### Functional annotation in 9q31

SNPs in LD (r^2^>0.8) with the lead SNP rs10979182 SNPs were examined for their intersection with ENCODE data for histone marks, transcription factor binding sites and DNase hypersensitivity. RegulomeDB v1.1 (Boyle et al., 2012) was used to rank the SNPs based on likely regulatory function. The SNPs were also assessed using Haploreg v4.1 (Ward & Kellis, 2012), which includes expression quantitative trait loci (eQTL) data from multiple studies including GTEx (Lonsdale et al., 2013) and GEUVADIS (Lappalainen et al., 2013).

### 3C-qPCR in 9q31

3C libraries were generated in biological triplicate as previously described (Naumova, Smith, Zhan, & Dekker, 2012). 20-30 million crosslinked cells were lysed, digested, ligated and purified as described in the first section of the Capture Hi-C protocol above, omitting the biotinylation step. Control libraries were constructed using bacterial artificial chromosome (BAC) clones as described by (Naumova et al., 2012). A set of 11 minimally-overlapping BAC sequences was compiled to span the locus (chr9:110168556-111889073, hg19): RP11-795I4, CTD-2258L2, RP11-762G1, RP11-358A7, RP11-779J13, CTD-2517A7, RP11-454G15, RP11-585H18, CTD-2333H8, CTD-2649N21 and RP11-316H9. BAC clone identity was confirmed by PCR to amplify selected regions at either end of the sequence. BAC DNA was combined in equimolar quantities to a total of 20 µg and digested with HindIII overnight at 37°C. The DNA was purified with phenol-chloroform and precipitated in ethanol for several hours at -20°C. Ligation was performed at 16°C overnight using T4 DNA ligase. Two further phenol-chloroform extractions were performed followed by a chloroform extraction and the DNA was precipitated in ethanol for several hours at -20°C. 3C libraries and BAC control libraries were quantified using a Qubit dsDNA BR kit.

qPCR was carried out using SYBR Green or TaqMan technology to determine interaction frequencies in the 9q31 locus. Unidirectional primers were designed using Primer3 (http://primer3.ut.ee/) (Untergasser et al., 2012) to complement sequences approximately 50-100 bp from the target HindIII cut site (tables 1-2). For the TaqMan experiment, an additional TaqMan probe was designed to bind to a region between the anchor primer and the restriction cut site. For each anchor fragment, a primer was designed to target a short-range control region located 2 – 3 fragments further along the sequence. qPCRs were carried out in technical triplicate and 10-fold dilutions of the BAC template (50 – 0.005 ng) were included alongside the 3C library templates for each tested interaction. For each primer pair, BAC curves were generated from Log_10_ of the library concentration against the average Ct value across PCR triplicates. The relative interaction frequencies were calculated in the following manner:

**Interaction Frequency (F) = 10^((Ct – i)/s)^**; where

**Ct** = Measured cycle threshold value of 3C library (mean of PCR triplicates)
**i** = Y intercept of BAC curve
**S** = Slope of BAC curve

The interaction frequencies were then normalised to the short-range control as follows:

**Relative Interaction Frequency (R) =** $\frac{\boldsymbol{F}[\boldsymbol{short}-\boldsymbol{rangecontrol}]}{\boldsymbol{F}[\boldsymbol{testinteraction}]}$; where **F** = Interaction frequency

Significant interactions were detected using one-way ANOVA in GraphPad Prism 7 with Dunnett’s or Tukey’s test for multiple comparisons dependent on whether there was a single negative control region or each interaction was compared to all other interactions, respectively.

Table 1. Fragments targeted in the first 9q31 (*KLF4*) 3C-qPCR assay

| Fragment name | Fragment designation | HindIII fragment location (hg19) | Primer sequence |
| --- | --- | --- | --- |
| Ps enhancer 3 (rs6477612) | Anchor | chr9:110810596-110816598 | CAGGGTGTCTAGGAGGTCTTC |
| Enhancer 3 SR | Short-range control | chr9:110824104-110824379 | ACCACATTCCTCTTTTAGCCC |
| RP11-363D24.1 | Target | chr9:110195017-110201199 | AGGAGCTGCATGATTCACCA |
| *KLF4* Centromeric 1 | Target | chr9:110237781-110238441 | GCAGGGCAATGGTCAATTCC |
| *KLF4* Centromeric 2 | Target | chr9:110241992-110244654 | CACACCTAGGAGCCCACAG |
| *KLF4* gene and promoter | Target | chr9:110244654-110255868 | TGCTTTGAAATGAAATCCCTGC |
| Intergenic A | Target | chr9:110452475-110459966 | GGCCAGCTAAGATTCACTGC |
| Intergenic C | Target | chr9:111343493-111346822 | TGAGTACACGCATCTTTTCCT |
| *IKBKAP* & *FAM206A* promoter | Target | chr9:111692307-111703976 | AGCTGTGGTCAATTGGCATT |
| *CTNNAL1* promoter | Target | chr9:111770041-111776135 | GGTGGCGAGGAGAAACAAAA |

Table 2. Fragments targeted in the second 9q31 (*KLF4*) 3C-qPCR assay

| Fragment name | Fragment designation | HindIII fragment location | Primer sequence |
| --- | --- | --- | --- |
| *KLF4* gene and promoter | Anchor | chr9:110244654-110255868 | TGCTTTGAAATGAAATCCCTGC |
| *KLF4* gene and promoter | TaqMan probe | chr9:110244654-110255868 | GCTTGCAGCTTTCACAAGGT |
| *KLF4* gene body SR | Short-range control | chr9:110257072-110259991 | GCAAACTCCTCTTATATCCAGGG |
| Intergenic A | Target | chr9:110452475-110459966 | GGCCAGCTAAGATTCACTGC |
| rs10979182 LD region 1 | Target | chr9:110765872-110770045 | TCGAAGACAGGTTGTTGGGA |
| rs10979182 LD region 2 | Target | chr9:110772357-110775650 | CTAAGGCCTGCAATGAAGACA |
| rs10979182 LD region 3 | Target | chr9:110780164-110782236 | GAAGAAGCATCCCATGGCTG |
| rs10979182 LD region 4 | Target | chr9:110783715-110788661 | TAAACCCAAGACAGTGCTGC |
| rs10979182 LD region 5 | Target | chr9:110793130-110796043 | GGCACCTGAGGCATACAATG |
| rs10979182 LD region 6 | Target | chr9:110798743-110801016 | AAATCATGTCCTTGGTGCCC |
| rs10979182 LD region 7 | Target | chr9:110801022-110808470 | TGGCTACATCCAGAGTTGCT |
| rs10979182 LD region 8 | Target | chr9:110810596-110816597 | TCTTCGCTTCCTGTGGGC |
| rs10979182 LD region 9 | Target | chr9:110816603-110821888 | TCAGCTCTTCAAGTTCTCATTCT |
| rs10979182 LD region 10 | Target | chr9:110824384-110829060 | AACCAAGTCACAGAGAAGGCT |
| rs10979182 LD region 11 | Target | chr9:110833319-110838267 | CTAATTGCTGCAGGACCCAC |
| Intergenic B | Target | chr9:110930759-110940017 | CTTGGAGTAGCTTGCTGAGG |
| Positive 1 | Target | chr9:111035783-111036598 | CTCCTCCAATCCCAAGCTGA |
| Positive 2 | Target | chr9:111038412-111038716 | CAGACAAAGACTGGACCCCA |
| Intergenic C | Target | chr9:111343493-111346822 | TGAGTACACGCATCTTTTCCT |

### ChIP-qPCR in 9q31

Chromatin immunoprecipitation (ChIP) libraries were generated as previously described (McGovern et al., 2016). Briefly, 10 million cross-linked cells were lysed and the chromatin was fragmented to optimal lengths of 200-400 bp using a Covaris S220. A volume of chromatin corresponding to approximately 1 million cells was immunoprecipitated with a rabbit polyclonal antibody for H3K4me1 (Abcam ab8895) or H3K27ac (ab4729) or a negative control rabbit IgG (Diagenode C15410206) with a mixture of Dynabeads A and G at 4°C overnight. After washes, the DNA was eluted with proteinase K at 62°C for 2 hours and then 95°C for 10 minutes. The DNA was purified using Purelink Quick PCR purification columns (Life Technologies). ChIP enrichment was measured at loci of interest by qPCR using SYBR Green or TaqMan technology. The data were normalised by calculating the percentage of total chromatin that was immunoprecipitated in comparison with an input sample. Negative controls included no-antibody and IgG precipitated samples.

For SYBR experiments, primers were designed using Primer3 (http://primer3.ut.ee/) to target regions of 100-200 bp encompassing likely regulatory SNPs or enhancers in 9q31 (Table 3). For the allele-specific analysis at rs6477612, a TaqMan SNP genotyping probe was acquired that detected C (risk) or T (protective) alleles (Applied Biosystems, assay ID C__29343482_10). The difference in antibody binding to each allele was determined by the percentage of ChIP enrichment in comparison with the signal for each allele obtained from the input sample. ChIP-qPCR data were analysed in GraphPad Prism using two-way ANOVA. Experiments were performed in biological triplicate (HaCaT and My-La lines) or duplicate (NHEK cells).

Table 3. Primers used in ChIP experiments

| Target | Primer name | Sequence |
| --- | --- | --- |
| 9q31 putative enhancer 1 | 27ac_ChIP_Pool1F1 | CAGAAGCTGTGAGAGGCTGA |
|  | 27ac_ChIP_Pool1R1 | ACTGAGGCAGCCGTAATGTT |
| 9q31 putative enhancer 2 | 27ac_ChIP_Pool2F1 | GAGCTGGGGAATGTGTGTTT |
|  | 27ac_ChIP_Pool2R1 | AAATGTGTTTGCCCTGGAAG |
| 9q31 putative enhancer 3 | 27ac_ChIP_Pool3F1 | ATCCCTTGTTTAGGGCTTGG |
|  | 27ac_ChIP_Pool3R1 | TGGACCTAGGCTTGCCTCTA |
| 9q31 putative enhancer 4 | 27ac_ChIP_Pool4F1 | TGACTGACCCAAGGTCACAT |
|  | 27ac_ChIP_Pool4R1 | GCATATACGGTTTCGGTGTG |
| *KLF4* promoter | CHIP_KLF4_PROM_F1 | CCTGAACCCCAAAGTCAACG |
|  | CHIP_KLF4_PROM_R1 | CGGACCTACTTACTCGCCTT |

### CRISPR activation in 9q31

CRISPR activation using the catalytically inactive Cas9 (dCas9)-P300 complex was performed in HaCaT cells to determine the role of the four putative enhancers in 9q31. Firstly, a HaCaT cell line stably expressing dCas9-P300 was generated using the CRISPRa plasmid pLV_dCas9-p300-P2A-PuroR plasmid (Addgene 83889) (Klann et al., 2017). Briefly, plasmid DNA was combined with third generation viral packaging components and polyethyenimine (PEI) in the ratio 1:6 (total DNA:PEI). The DNA:PEI complexes were added to Lenti-X 293T cells, which were then incubated for 72 hours, after which the lentivirus-containing supernatant was harvested and filtered to remove cell debris. Lentiviral transduction of 300,000 HaCaT cells was carried out using 1 mL of the un-concentrated lentivirus and 8 µg/mL polybrene. The cells were grown for several days before selection with 1 µg/mL puromycin for 7 days, after which the HaCaT dCas9-P300 cell population was maintained with 0.5 µg/mL puromycin.

To select sgRNA in 9q31, SNPs in r^2^> 0.8 with rs10979182 were prioritised by their overlap with enhancer elements, defined by active regulatory regions in NHEK according to ChromHMM (Ernst et al., 2011). sgRNA sequences were designed using the online CRISPOR tool (Concordet & Haeussler, 2018) to target loci within 200 bp of the prioritised SNPs (mean = 85 bp). sgRNA were selected based on specificity score and proximity to the SNP. In total there were 27 SNPs; two of these could not be targeted by sgRNA within 200 bp (rs7019552 and rs11355519) and another two SNPs, rs4979624 and rs7029094, were captured by a single sgRNA targeting the intervening region. In total, therefore, there were 24 sgRNA; these were grouped into four pools of 5-7 SNPs to target the four putative enhancers (Table 4; Figure 1). The sgRNA sequences were cloned into the pLKO5.sgRNA.EFS.GFP plasmid (Addgene 57822) (Heckl et al., 2014) and equimolar plasmid pools were generated for each enhancer. The plasmid pools were then packaged using the same lentiviral method as the dCas9-P300 plasmid. The guide pools were introduced into the stable HaCaT dCas9-P300 cells using a second round of lentiviral transduction and cells that had integrated the sgRNA plasmids were isolated by flow cytometry for GFP. RNA was extracted from the sorted cells using the RNeasy mini kit (Qiagen). qPCR was performed to assay gene expression using the TaqMan RNA-to-Ct 1-step kit (Thermofisher Scientific) using the following TaqMan assays for genes in the 9q31 locus: *RAD23B* (Hs00234102_m1), *KLF4* (Hs00358836_m1), *ACTL7A* (Hs00246418_s1), *ACTL7B* (Hs00246411_s1), *IKBKAP* (Hs00175353_m1), *FAM206A* (Hs00607423_m1) and *CTNNAL1* (Hs00972098_m1). Delta-delta Ct analysis was conducted against a control HaCaT dCas9-P300 cell line transduced with the sgRNA plasmid containing a previously-published scrambled, non-targeting insert (Scramble2; (Lawhorn, Ferreira, & Wang, 2014)). Two housekeeping gene assays, *TBP* (Hs00427620_m1) and *YWHAZ* (Hs01122445_g1), were used for normalisation. For the CRISPRa Pool with the greatest impact on *KLF4* expression, RNA-seq and gene set enrichment analyses were performed as described above. In addition, differentially expressed genes were processed through the STRING database to identify potential protein-protein interaction networks (Szklarczyk et al., 2015)

Table 4. sgRNA used in the CRISPRa experiment

| Purpose | SNP/gene | Protospacer Sequence |
| --- | --- | --- |
| Scrambled non-targeting control (see Lawhorn et al., 2014) | N/A | AACAGTCGCGTTTGCGACT |
| IL1RN positive control (Perez-Pinera et al., 2013) | IL1RN promoter | CATCAAGTCAGCCATCAGC |
| SLC4A1 positive control (Weissman Lab) | SLC4A1 promoter | GTCAGGAGAACCATGGGGACC |
| 9q31 Pool 1 | rs10816609 | CTAATAAGCATCATCGCCCA |
|  | rs35078320 | GTCTCTCTTTAGGCTATCGT |
|  | rs10816610 | GGCTCTTATTCATAGTGTTA |
|  | rs4979624/rs7029094 | AGCTCTTGATATGACCTCAA |
|  | rs10512368 | GCCCAAGACTATGGAATTGT |
|  | rs10816611 | GATAATAGATCTTCCTACAG |
| 9q31 Pool 2 | rs1361371 | AAAGTCTAGGTCTCGAATCC |
|  | rs10816617 | GTGAATGCTGATTGTAACCC |
|  | rs10816618 | ATTTATGTATACATCGATTG |
|  | rs10118193 | AGATTCTTGAGAGCGGTAGC |
|  | rs1914513 | CCACACTGCTGCATTGATTC |
|  | rs10217259 | AAGGGGCTAATGCCTGTTCA |
| 9q31 Pool 3 | rs10979180 | CCACCAGGGACCCAAAAGGT |
|  | rs113137157 | AATCCCTTGTTTAGGGCTTG |
|  | rs55975335 | CTGAAGGTCCTTTGTTATTG |
|  | rs6477612 | TGGTTTCGAGATTCCTAAAC |
|  | rs6477613 | TGCGTGAGGCTGTACATTAT |
| 9q31 Pool 4 | rs1318148 | GATTGGAGACTCGCCATCAG |
|  | rs892687 | CAATAAAAGCCGGGTAGACC |
|  | rs1369190 | AGCTCAGCCTGATTCTCATG |
|  | rs10979182 | GTCAGCCTAGAGGTTTCTAG |
|  | rs4978668 | CGGACTGGTACTTTAGGTGC |
|  | rs2417842 | GGCACAATGCCTTCGAAGGT |
|  | rs4978343 | TTCGTATATCAGTCTTTGCC |

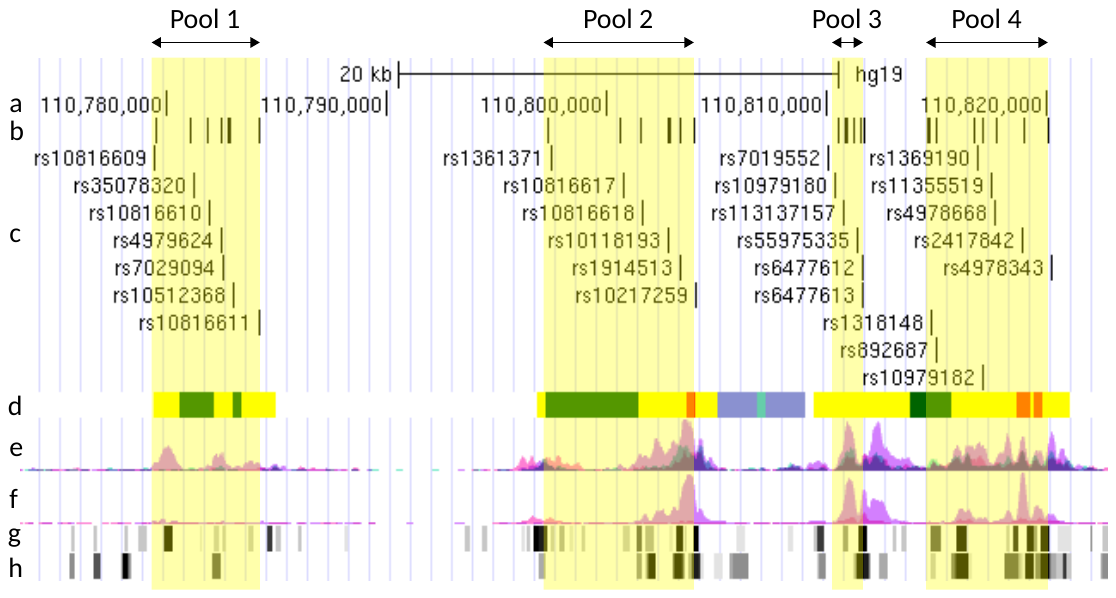

Figure 1: sgRNA pools targeting the four putative enhancers in the psoriasis susceptibility locus at 9q31. a) location on chromosome 9; b) sgRNA locations; c) SNPs in LD with rs10979182 overlapping putative enhancers; d) ChromHMM segments in NHEK where red, yellow and green indicate “active TSS”, “enhancers” and “transcription” respectively; e) H3K4me1 (ENCODE); f) H3K27ac (ENCODE); g) DNase clusters (ENCODE); h) transcription factor ChIP (ENCODE).

In addition to the above, the validity of the HaCaT dCas9-P300 system was first tested using sgRNA directed to the promoters of *IL1RN* or *SLC4A1*; these sgRNA have previously been shown to be effective in CRISPRa (IL1RN: sgRNA CR3 (Perez-Pinera et al., 2013); SLC4A1: Weissman Lab). The upregulation of these genes was detected by TaqMan qPCR for IL1RN (Hs00893626_m1) and SLC4A1 (Hs00978607_g1) (Figure 2).

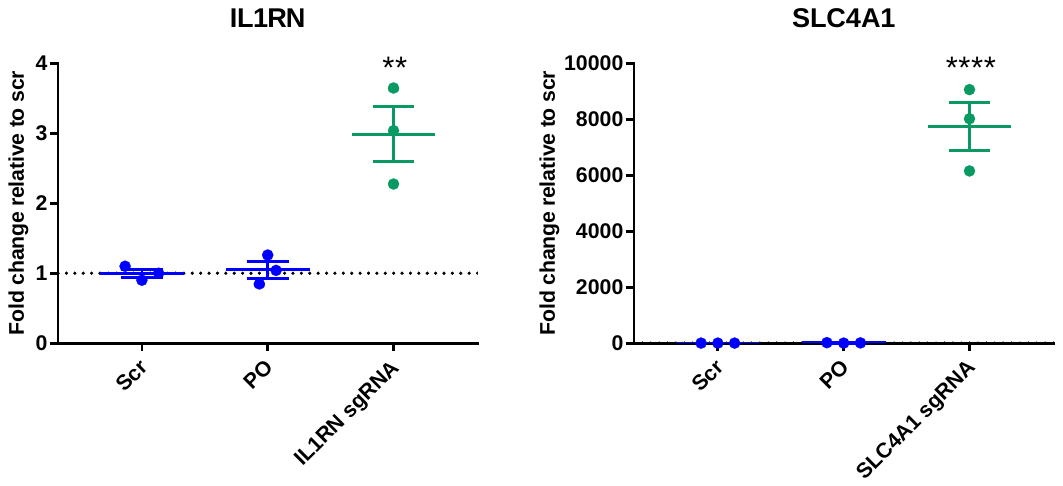

Figure 2: Effect of sgRNA targeting *IL1RN* and *SLC4A1* promoters in HaCaT dCas9 P300 cells. HaCaT cells expressing dCas9 P300 were transduced with sgRNA plasmids containing previously-published sgRNA for the *IL1RN* promoter or the *SLC4A1* promoter, in biological triplicate. Control cell lines were generated by transducing HaCaT dCas9 P300 cells with a sgRNA plasmid containing a scrambled guide (Scr) or no guide insert (Plasmid only; PO). qPCR was carried out using TaqMan assays for *IL1RN* or *SLC4A1*. Housekeeping genes used were *TBP* and *YWHAZ*. Graphs show fold change of gene expression relative to the cells containing the scrambled sgRNA. One-way ANOVA was carried out in GraphPad Prism: in both cases, the cells containing the targeting sgRNA had significantly higher gene expression than the scrambled control (P=0.002 for *IL1RN*; P=0.0001 for *SLC4A1*). Asterisks denote P < 0.05. Graphs show the mean fold-change in comparison with scrambled guide, +- SEM of triplicate cell lines.
