## supplementary results for "Chromatin-based techniques map DNA interaction landscapes in psoriasis susceptibility loci and highlight *KLF4* as a target gene in 9q31"

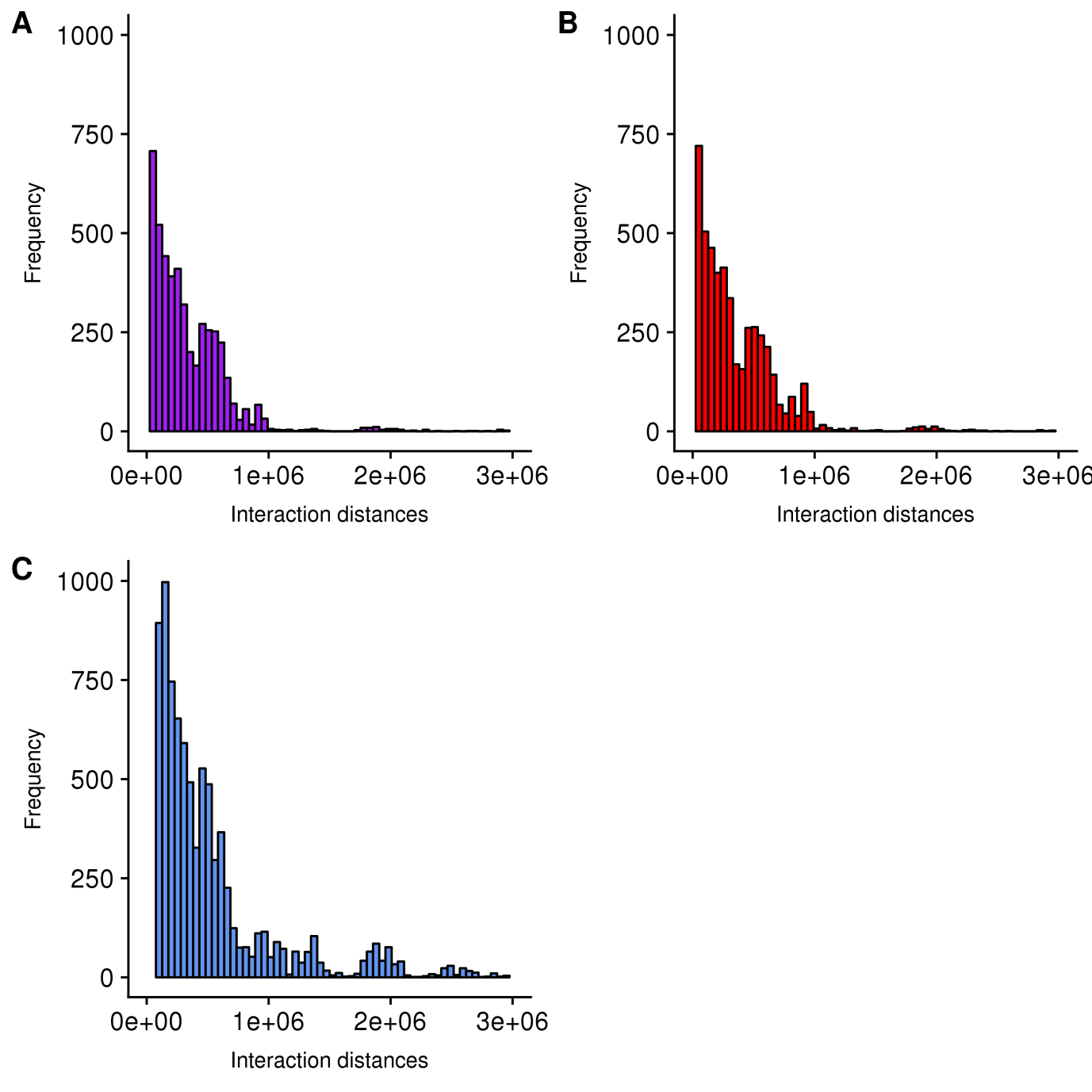

Figure S1: Frequency distributions of distances between psoriasis bait fragments and interacting fragments in the CHi-C experiment. The frequency of interactions is shown for 500 kb bins up to 3 Mb in HaCaT unstimulated (A), HaCaT stimulated (B) and My-La cells (C).

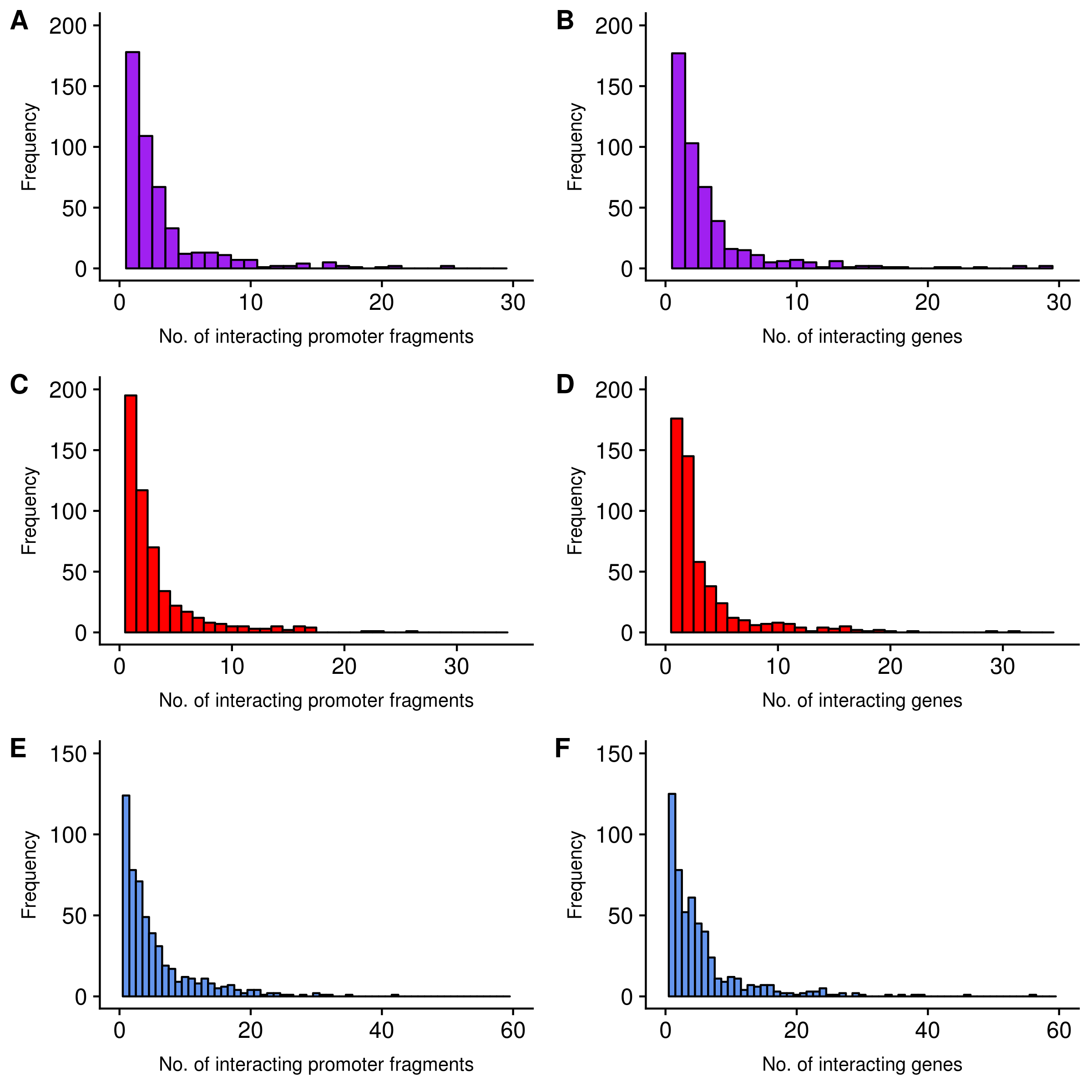

Figure S2. Frequency distributions of the number of interactions with promoter fragments per psoriasis-associated bait fragment. To determine the frequency distribution of psoriasis bait-promoter interactions, the data was firstly restricted to interactions between psoriasis-associated bait fragments and promoter fragments (“Promoter Interactions”). Next, the number of promoter fragments per bait fragment was counted. Of those promoter fragments, the number of corresponding gene promoters was determined. This was necessary because some gene promoters share the same fragment, and some gene promoters are found in more than one fragment. The number of interacting promoter fragments per bait fragment in Promoter Interactions are shown for HaCaT unstimulated (A), HaCaT stimulated (C) and My-La (E). The number of corresponding gene promoters are shown for HaCaT unstimulated (B), HaCaT stimulated (D) and My-La (F). The interaction frequencies are shown in bins of 1.

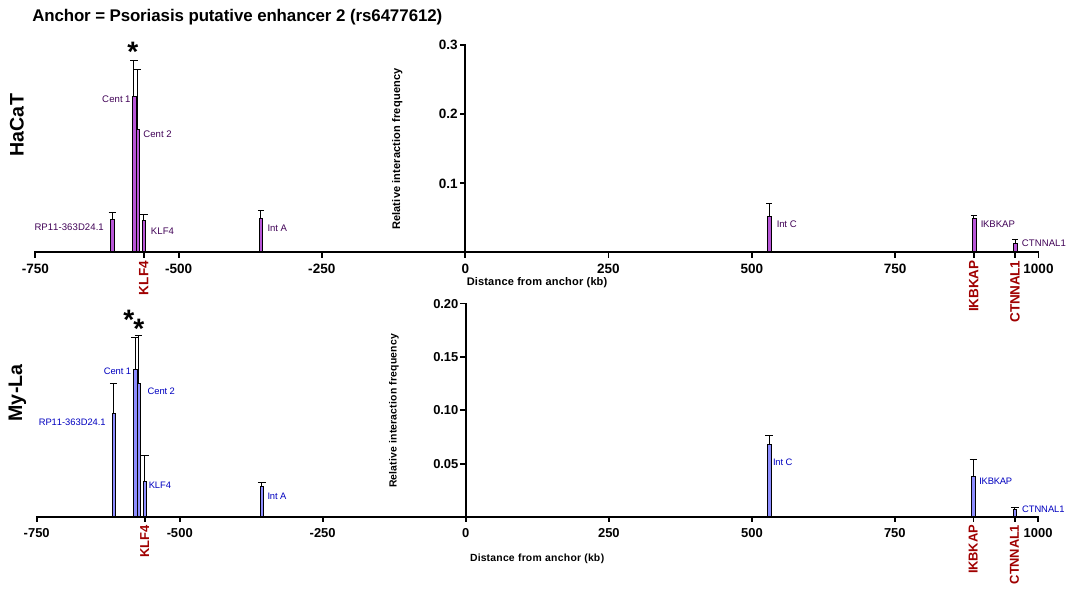

Figure S3. 3C-qPCR results in the 9q31 locus anchored at the HindIII fragment containing the third psoriasis-associated putative enhancer (rs6477612). qPCR was carried out on HaCaT and My-La 3C libraries using SYBR® Green as the reporter. The anchor fragment at the third psoriasis-associated enhancer is at distance 0 kb. Test fragments were selected in and around *KLF4*, two points in the gene desert and at fragments containing gene promoters for *IKBKAP*, *FAM206A* and *CTNNAL1*. Interactions were normalised to a short range control. Asterisks denote fragments that had a significantly higher relative interaction frequency than one or more of the other tested fragments, after multiple testing (one-way ANOVA, adjusted P-value < 0.05). Bars show mean + SEM of triplicate 3C libraries. Abbreviations: Cent, centromeric; Int, intergenic

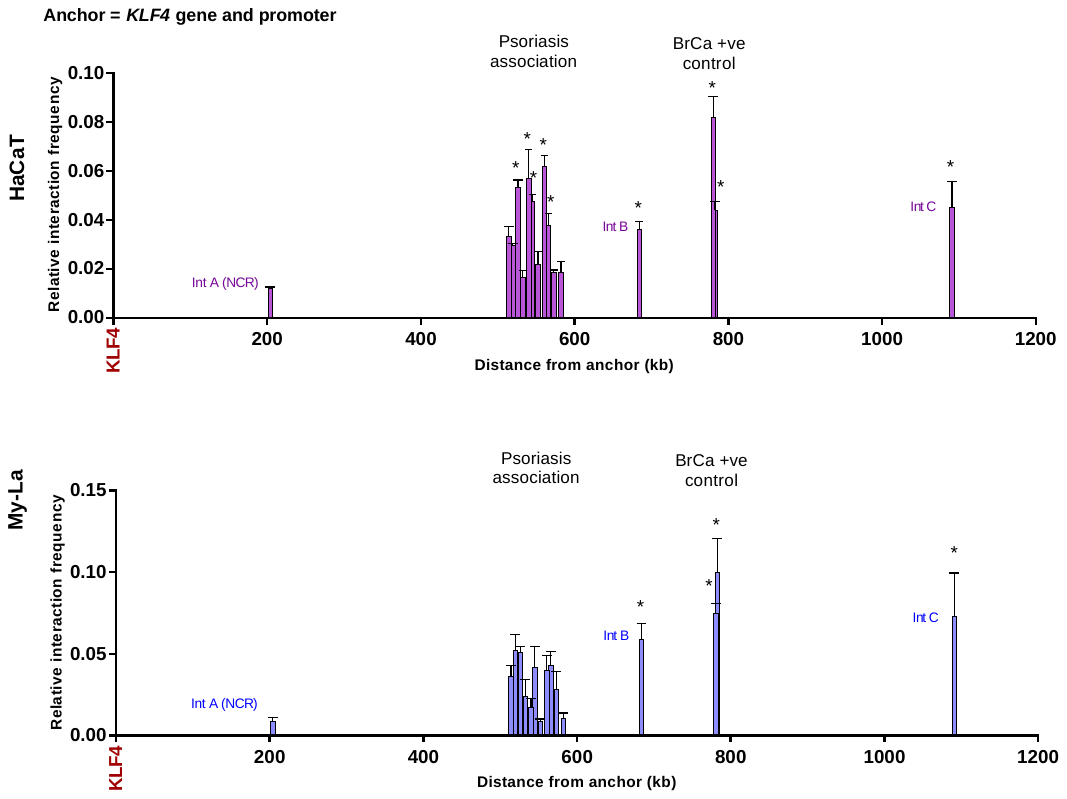

Figure S4. 3C-qPCR results in the 9q31 locus from the HindIII fragment containing the *KLF4* gene and promoter. qPCR was carried out on HaCaT and My-La 3C libraries using TaqMan® as the reporter. The anchor fragment (distance 0) contained the entire *KLF4* gene and promoter. An intergenic fragment located approximately 200 kb from the anchor fragment was utilised as a negative control region. Eleven test fragments were selected at regular intervals across the psoriasis association. The positive controls in the Dryden BrCa region were included. Asterisks denote fragments that had a significantly higher relative interaction frequency than the NCR (one-way ANOVA, adjusted P-value < 0.05). Bars show mean + SEM of triplicate 3C libraries. Abbreviations: Int, intergenic; NCR, negative control region; BrCa, breast cancer.

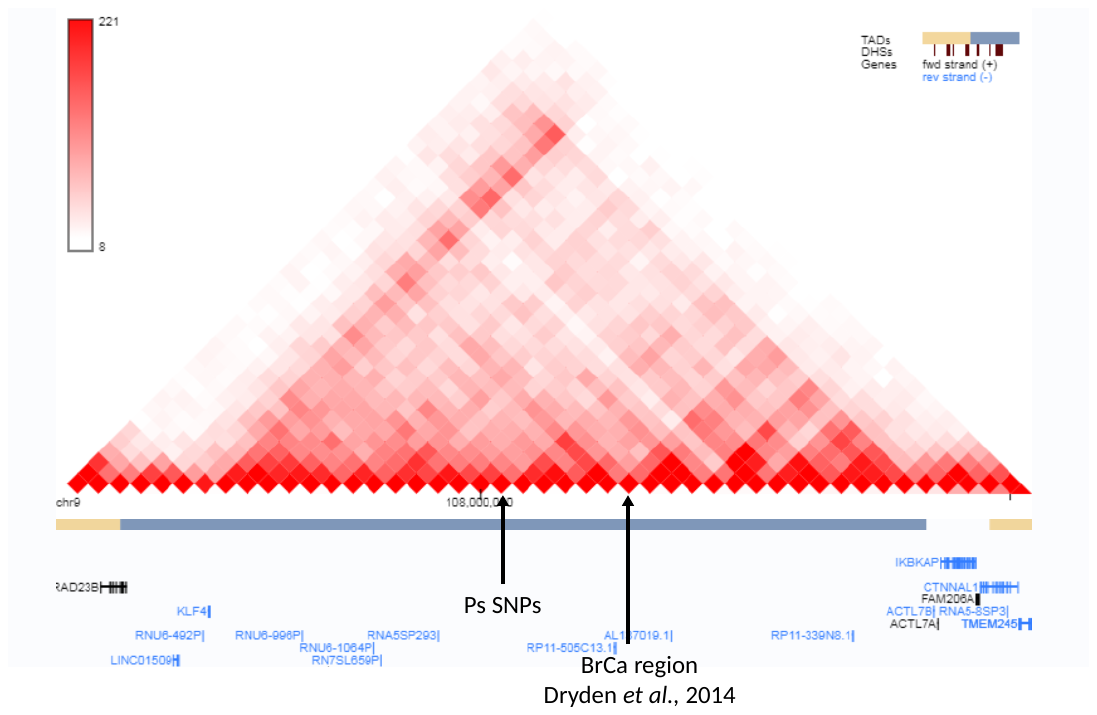

Figure S5. Previously reported HiC interaction data in NHEK cells in the 9q31 locus (Rao et al, 2014). Interactions are indicated between KLF4 and the gene desert, including the psoriasis SNPs and the breast cancer region shown in a CHiC experiment by Dryden et al (2014). Image created using the YUE lab 3D Genome Browser.
